# The Intermediate CA1 and CA3 Subfields of the Hippocampus Differentially Encode Social Recognition

**DOI:** 10.1101/2025.10.27.684580

**Authors:** Benjamin Dykstra, Aidan Lin, Paul Kim, Cem Sevinc, Gordon J. Berman, Malavika Murugan

## Abstract

Social recognition requires linking information about others to the spatial contexts in which they are encountered, a computation thought to engage the hippocampus. Although this process depends on coordinated activity across the hippocampal network, it remains unclear how specific subfields differentially contribute to social recognition. Of particular interest are the intermediate CA1 (iCA1) and CA3 (iCA3) subfields, which have both been shown to be causally involved in social recognition. Historically, the dense feed-forward projections from CA3 to CA1 led to the view that these subfields form a hierarchical circuit, with CA1 primarily integrating and refining computations from CA3. However, recent causal manipulations of the iCA1 and iCA3 have revealed distinct and separable contributions to approach and avoidance behavior, suggesting functional specialization across the subfields. This functional divergence suggests that iCA1 and iCA3 may differentially encode social recognition information. To test this possibility, we performed one-photon calcium imaging in freely behaving mice to record endogenous activity from the iCA1 and iCA3 during a novel behavioral paradigm that disentangles social identity from spatial position. Notably, we found that iCA3 neurons more robustly discriminated between novel and familiar conspecifics, while iCA1 neurons more strongly encoded spatial position. These results identify the iCA3 as a key locus for social novelty encoding and suggest distinct contributions of the iCA1 and iCA3 to social recognition.

## Introduction

Almost every social encounter an animal engages in begins with an assessment: am I interacting with someone I know or someone new? Recognizing familiar individuals allows animals to make informed behavioral decisions based on prior interactions with that individual. Equally important is the ability to detect novel conspecifics who may pose potential threats or present opportunities. In mice, the ability to discriminate between novel and familiar conspecifics is well established^1^, and depends on hippocampal circuitry^2,3^.

The rodent hippocampus is composed of several subfields that are thought to differentially contribute to social recognition. Among these subfields, the CA2 has been the most intensively studied^4–8^. Lesioning or inactivating the CA2 disrupts rodents’ natural preference for investigating novel conspecifics over familiar conspecifics^9,10^. Furthermore, electrophysiological recordings in the CA2 have identified neurons that encode the identity of novel and familiar conspecifics^11^. However, recent evidence suggests that social recognition relies on a broader hippocampal network. In particular, the intermediate CA1 (iCA1) and CA3 (iCA3) have emerged as important contributors. Inhibiting the intermediate and ventral CA1 and CA3 regions disrupt social novelty preference, whereas inhibition of the dorsal CA1 and CA3 does not^12,13^. Additionally, Okuyama et al., 2016 and Tao et al, 2022 found that ventral CA1 neurons encode social recognition information, with a larger proportion of neurons responding to familiar conspecifics over novel conspecifics ^12,14^. Yet no studies have directly recorded neuronal activity in the iCA3 during social behavior, leaving its contribution largely unknown. Given the strong monosynaptic projections from the iCA3 to the iCA1^15,16^, understanding how these two subfields interact is critical for revealing how social recognition signals emerge and are transformed within hippocampal circuits.

Furthermore, the iCA1 and iCA3 may implement distinct but complementary computations relevant to social recognition processing. Historically, the strong feed-forward projections from CA3 to CA1 led to the view that these subfields operate within a hierarchical circuit, encoding shared information at distinct representational levels^17^. While this hierarchical organization suggests complementary processing of related inputs, emerging evidence indicates that the subfields also make distinct behavioral contributions. Both regions send direct projections outside the hippocampus^18–20^, and recent causal manipulations of iCA1 and iCA3 projections to the lateral septum have been shown to differentially bias approach and avoidance behaviors^21^. Together, these findings suggest that, in addition to processing information at different levels of abstraction, iCA1 and iCA3 may also exhibit functional specialization. Whether this divergence extends to encoding social recognition information remains unknown.

A major obstacle in addressing these questions is the strong spatial encoding observed in both the iCA1 and iCA3^22^, potentially masking or modulating more subtle social signals. In the most widely used behavioral assay to study social recognition and memory, the social discrimination test (SDT), different conspecifics are presented at distinct spatial locations.^12,23^ While the SDT has provided valuable insight into social recognition, the spatial constraints of the assay make it difficult to disentangle social from spatial encoding. This challenge is amplified when studying the hippocampus, which is thought to contextualize diverse forms of information, including social representations, into a spatial framework^22,24^. Thus, location of the conspecific can often serve as a proxy for social identity. These limitations highlight the need for new behavioral paradigms that preserve naturalistic social interactions while minimizing social–spatial confounds.

Here, we used one-photon cellular-resolution calcium imaging to record the endogenous activity of iCA1 and iCA3 neurons in two separate cohorts of mice. Mice participated in both a standard SDT and a novel paradigm, the Linear Presentation Assay (LPA), which aims to disassociate social from spatial encoding. In the SDT, we found that social recognition representations in both the iCA1 and iCA3 subfields were conjunctively encoded with spatial information. In the LPA, an automated assay where multiple conspecifics are randomly presented at the same spatial location, we found that both iCA1 and iCA3 neurons carry purely social information, but that iCA3 neurons more distinctly discriminate between novel and familiar conspecifics relative to the iCA1. Longitudinal tracking further revealed that, unlike spatial representations, social novelty representations did not persist across days, suggesting that the iCA3 novel/familiar signals may support real-time social discrimination rather than a general signal of social novelty. These results identify the iCA3 as a key site for social novelty encoding and reveal a functional dissociation of roles between the iCA1 and iCA3. Together, they support a broader view of the hippocampus as a multiplexed system that segregates social information into distinct representations across subfields.

## Results

### Cellular Resolution Imaging of iCA1 and iCA3 During Social Discrimination Test

To determine whether the iCA3 encodes social recognition information and how these representations compare to those in the iCA1, we performed one-photon cellular resolution calcium imaging in two separate cohorts of mice. We expressed the calcium indicator GCaMP6f in the intermediate hippocampus and implanted a gradient-index (GRIN) lens targeting either the iCA1 (n =15 male mice) or the iCA3 (n = 12 male mice; Fig. 1A). For each subfield, we used two different lens targeting strategies, straight lenses (iCA1 n = 5; iCA3 n = 1) and prism lenses (iCA1 n = 10; iCA3 n = 11; Suppl. Fig. 1D, E). Prism lenses were implanted posterior to the target region, enabling imaging from coronal planes (Fig. 1B; Suppl. Fig. 1B, C), and straight lenses were implanted dorsally to capture horizontal planes (Suppl. Fig. 1A). Because this approach produced distinct lens trajectories, it mitigates the likelihood of consistent off-target damage across animals. GRIN lens placement and viral expression were verified post hoc by histological analysis. In total, we recorded 1506 neurons from 15 mice in the iCA1 and 1017 neurons from 12 mice in the iCA3 (Fig. 1D).

**Figure 1.**
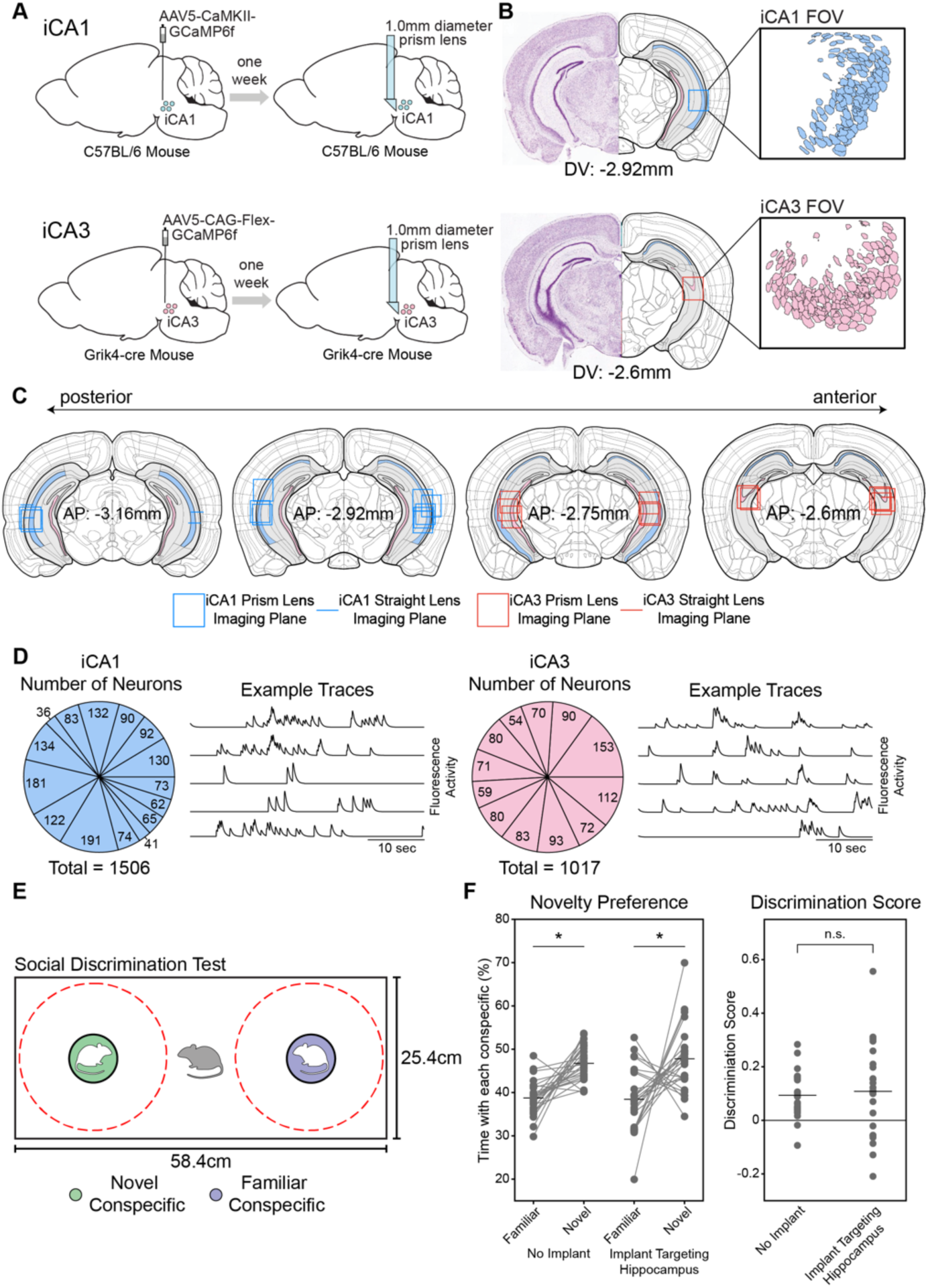
Recording Cellular Resolution Activity in the iCA1 and iCA3. **A)** Schematic of viral injection of GCaMP (left) and GRIN lens implantation (right) targeting iCA1 (top panels) and iCA3 (bottom panels). **B)** Representative coronal sections showing imaging planes (left) and the corresponding fields of view (FOVs) from two example mice: iCA1 (top) and iCA3 (bottom). Anatomical sections were adapted from the Allen Brain Atlas. **C)** Reconstruction of imaging planes across all iCA1 and iCA3 mice. **D)** Average number of neurons collected per mouse in iCA1 (left) and iCA3 (right) across imaging sessions. Example calcium traces of individual neurons from representative animals are shown to the right of each pie chart. **E)** Schematic of the social discrimination test (SDT) behavioral assay used to assess novelty preference. **F)** (left) Mice preferentially investigated novel conspecifics over familiar conspecifics during the SDT in lens implanted mice and a no lens control cohort (lens implanted mice: paired-test (t(24) = 3.173, p = 0.004, n = 25; no lens: paired t-test t(25) = 5.674, p < 0.001, n = 26). (right) Lens implanted mice and no lens control mice did not differ in their discrimination score (t-test, t(49) = 0.398, p = 0.692).

Throughout this manuscript, we refer to our recording site as the intermediate hippocampus to be consistent with anatomical and molecular boundaries that divide the rodent hippocampus along the dorsoventral axis^25^. However, the terms “intermediate”^26–28^ and “ventral”^12,29–31^ hippocampus are applied inconsistently across the literature, and studies using these terms often show substantial coordinate overlap. Given known functional gradients along the dorsoventral axis^12,13^, we have provided our stereotaxic coordinates (Methods), histological verification (Suppl. Fig. 1), and reconstructions of lens placement (Fig. 1C) to encourage comparisons with other studies that may use different naming conventions or anatomical boundaries.

We confirmed that the lens-implanted imaging mice retained the ability to discriminate between novel and familiar conspecifics. Imaging mice were run on the Social Discrimination Test (SDT) in which mice freely investigated a novel and a familiar conspecific confined at opposite ends of a behavioral arena (Fig. 1E). Imaging cohorts spent significantly more time investigating a novel conspecific than a familiar conspecific, similar to a control cohort without implants (Fig. 1F; statistics reported in figure legend).

We also conducted a free interaction test in which imaging mice were placed into a cylindrical arena with either a novel or a familiar conspecific (Suppl. Fig. 2A). This assay captures naturalistic social interactions beyond the structured choices measured during more constrained tests. Behavioral recordings were annotated using a supervised SimBA network^32^ trained to detect social investigation (Suppl. Fig. 2B, C). In this assay, imaging mice showed a significant preference for investigating the novel mouse over the familiar mouse (Suppl. Fig. 2D; statistics reported in figure legend), matching the behavior of a control cohort lacking implants. Together, these findings demonstrate that imaging mice retained the ability to discriminate between novel and familiar individuals, indicating that the surgical procedures and implants did not compromise their ability to preferentially investigate novel over familiar conspecifics.

Next, we recorded neural activity from the iCA1 and iCA3 while imaging mice investigated two conspecifics confined to cages placed in opposite corners of a large SDT behavioral arena (70 × 70 cm). The large SDT arena was chosen to account for the larger place fields sizes in more ventral portions of the hippocampus^33,34^, reducing potential overlap in place fields centered on either corner. Additionally, in this arena, it took several seconds for the imaging mouse to run from one conspecific to the other, thereby reducing the chance that neural activity from one interaction would carry over into the next (see Movie 1).

The imaging mice were allowed to explore the SDT arena in two different behavioral conditions to probe different aspects of social encoding (Fig. 2A). In the first condition, we recorded activity while mice investigated two novel conspecifics to determine whether iCA1 and iCA3 neurons encode the identity of individual mice. In the second condition, we recorded activity while the imaging mice investigated one novel and one familiar conspecific (a same-sex cagemate of the imaging mouse) to determine if neurons encode social novelty. Because the CA1 and CA3 regions of the hippocampus encode spatial information^22^, we swapped the positions of the conspecifics midway through a 40-minute behavioral recording to dissociate social and spatial information (Fig. 2B).

**Figure 2.**
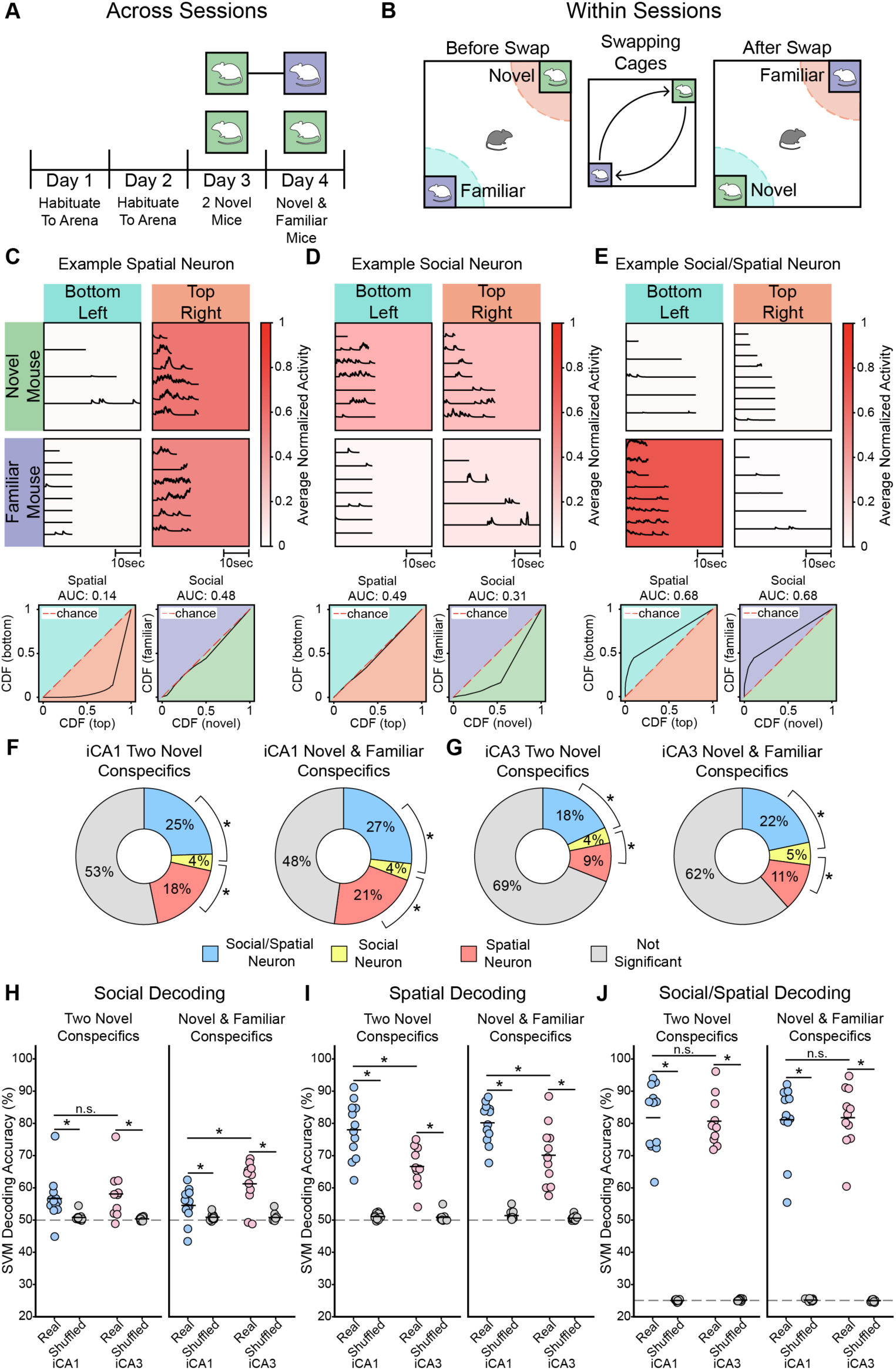
Social and Spatial Encoding in the iCA1 and iCA3. **A)** Timeline of behavioral experiments in the SDT assay. **B)** Schematic of the SDT showing the placement of conspecifics before and after the swap. **C)** Trial-by-trial responses of an example neuron during investigation of novel and familiar conspecifics in two distinct spatial locations. Trials organized in ascending order of time spent in the social zone in each condition. Boxes are color coded by the average response across trials with dark red representing high activity and white representing low activity. Below, receiver operating characteristic (ROC) curves and corresponding area under the curve (AUC) for spatial (left) and social (right) discriminability of the example neuron shown above, computed by comparing the cumulative distribution of neural activity across conditions (see Methods). This example neuron is spatially modulated but not socially modulated. **D)** Example neuron that is socially modulated and not spatially modulated. **E)** An example social–spatial neuron. This neuron preferentially fires in the bottom left but only in the presence of the familiar conspecific. **F)** Proportion of iCA1 neurons that display significant social– spatial preferences, neurons that only display a significant social preference and neurons that only display a significant spatial preference. Neural preferences for the 2-novel session are shown on the left and neural preferences in the novel-familiar conspecific session are shown on the right. In both conditions, there is a significantly larger proportion of spatial than social neurons (2-novel: z = 12.857, p < 0.001; novel–familiar: z = 13.876, p < 0.001) and a significantly larger proportion of social–spatial neurons than social neurons (2-novel: z = 16.523, p < 0.001; novel– familiar: z = 16.770, p < 0.001). **G)**, Same as (F), but for the iCA3. In both conditions there is a significant difference in the proportion of social and spatial neurons (2-novel: z = 4.499, p < 0.001; novel–familiar: z = 4.128, p < 0.001) and there is a significantly larger proportion of social–spatial neurons than social neurons (2-novel: z = 9.318, p < 0.001; novel–familiar: z = 9.178, p < 0.001). **H)** SVM decoding accuracy in the iCA1 (blue), iCA3 (pink) and shuffled traces (gray) when decoding conspecific identity from population neural data. Each point represents the average decoding accuracy of one mouse. Decoding accuracy was significantly above chance in both the two-novel condition (iCA1: Δ = 5.87%, p < 0.001, d = 4.07; iCA3: Δ = 7.69%, p < 0.001, d = 4.68) and the novel/familiar condition (iCA1: Δ = 3.67%, p < 0.001, d = 3.45; iCA3: Δ = 10.44%, p < 0.001, d = 7.22), with iCA3 decoding accuracy significantly higher than iCA1 in the novel/familiar condition, but not the two-novel condition (2-novel: Δ = 1.40%, p = 0.67, d = 0.65; novel/familiar: Δ = 6.72%, p = 0.009, d = 3.85). **(I)** Same as (H), but decoding whether the imaging mouse was in the bottom-left vs. top-right corner of the arena. Decoding accuracy was significantly above chance in both the two-novel condition (iCA1: Δ = 26.93%, p < 0.001, d = 15.56; iCA3: Δ = 15.79%, p < 0.001, d = 11.28) and the novel/familiar condition (iCA1: Δ = 28.82%, p < 0.001, d = 22.64; iCA3: Δ = 19.57%, p < 0.001, d = 9.93), with decoding accuracy significantly higher in the iCA1 than the iCA3 in both conditions (2-novel: Δ = 11.37%, p < 0.001, d = 5.13; novel/familiar: Δ = 10.03%, p = 0.002, d = 4.35). **(J)** Same as (H,I), using a multi-class SVM decoder to decode all four social-spatial conditions. Decoding accuracy was significantly above chance in both the two-novel condition (iCA1: Δ = 56.8%, p < 0.001, d = 28.09; iCA3: Δ = 55.5%, p < 0.001, d = 33.2) and the novel/familiar condition (iCA1: Δ = 56.03%, p < 0.001, d = 26.18; iCA3: Δ = 56.83%, p < 0.001, d = 29.23), with no significant difference in decoding accuracy between subfields (2-novel: Δ = 0.63%, p = 0.90, d = 0.23; novel/familiar: Δ = 0.63%, p = 0.91, d = 0.22).

Next, we classified neurons as spatially selective, socially selective, or selective for a specific social–spatial combination using a receiver operating characteristic (ROC) analysis (Fig. 2C-E). In both the iCA1 and iCA3, the proportion of spatially selective neurons was significantly greater than the proportion of socially selective neurons in both the two-novel conspecific condition and the novel/familiar condition (Fig. 2F,G; statistics reported in figure legend). Similarly, the proportion of conjunctive social–spatial neurons was significantly greater than the proportion of socially selective neurons in both conditions (Fig. 2F,G). Together, these findings indicate that iCA1 and iCA3 neurons were more likely to represent purely spatial information or a conjunction of social–spatial information, rather than purely social responses in the SDT. In addition to the pooled analysis across mice (Fig. 2F,G), these trends held true when examining the proportions of social, spatial, and social–spatial neurons across individual mice (Suppl. Fig. 3A). Furthermore, distributions of AUC values revealed a range of stimulus response strengths within each category (Suppl. Fig. 3B, C).

In both the iCA1 and iCA3, neurons with purely social preferences were evenly distributed between those preferring each conspecific (Suppl. Fig. 3D). We found iCA1 and iCA3 neurons with social–spatial selectivity for all four social–spatial combinations, with a larger fraction preferring conditions before the conspecific swap (Suppl. Fig. 3E).

Next, we used support vector machines (SVMs) to decode conspecific identity from population activity in the iCA1 and iCA3 (Suppl. Fig. 4). In both the two novel conspecific and novel/familiar conspecific conditions, both the iCA1 and iCA3 could decode conspecifics above chance levels (Fig. 2H). In the two novel conspecific condition, there was no significant difference between iCA1 and iCA3 decoding. However, in the novel/familiar condition, the iCA3 was able to decode conspecifics with significantly higher decoding accuracy compared to the iCA1 (Fig. 2H). We next tested whether spatial location could be accurately decoded from the same neural populations. SVMs significantly decoded whether the imaging mouse was located in the top right or bottom left corner in both conditions in both the iCA1 and iCA3, with the iCA1 having significantly higher decoding accuracy than the iCA3 in both conditions (Fig. 2I). Finally, we used multi-class SVMs to decode all four social–spatial combinations simultaneously. In both the two-novel and novel-familiar conditions, decoding was significantly above chance in both the iCA1 and the iCA3 (Fig. 2J).

We next compared decoding accuracy across social, spatial, and social–spatial decoders by normalizing the accuracy relative to chance levels to allow for direct comparisons (Suppl. Fig. 5A). Across both iCA1 and iCA3 cohorts, spatial decoding was significantly higher than social decoding and social–spatial decoding was significantly higher than social decoding and spatial decoding (Suppl. Fig. 5A).

To assess how decoding performance scaled with population size, we repeated each decoding analysis using increasing subsets of randomly sampled neurons within each mouse. For social, spatial, and social–spatial decoding, accuracy increased with neuron count and approached saturation (Suppl. Fig. 5B, C), implying that the sampled populations were large enough to capture most of the decodable information in both subfields.

Given that many neurons preferentially fired for specific social–spatial combinations, we next asked whether social and spatial representations at the population level remained stable across the entire session or whether they were similarly influenced by the conspecific swap. By comparing classifier confidence (distance to the decision boundary) and accuracy across time bins (Suppl. Fig. 5D, F), we found no consistent differences between pre-swap and post-swap decoding for either social or spatial classifiers (Suppl. Fig. 5E, G), suggesting that decoding performance was stable across an imaging session.

Together, these results demonstrate that in the SDT, in both the iCA1 and iCA3, social recognition representations are heavily influenced by spatial information. To better dissociate social and spatial representations, we next developed a novel behavioral assay designed to more directly test encoding of social identity independent of spatial location.

### Linear Presentation Assay, a Novel Social Recognition Behavioral Assay that Better Dissociates Social from Spatial Encoding

To investigate social recognition while rigorously controlling for spatial coding, we developed a novel behavioral assay: the Linear Presentation Assay (LPA). In this paradigm, an imaging mouse interacts with two conspecifics that are presented at the same spatial location. The assay consists of a long, narrow behavioral arena that the imaging mouse can freely explore. At one end of the arena is a barred interaction window. The stimulus cage, positioned behind the interaction window, sits on a programmable and automated conveyor belt that allows for multiple stimulus cages to be sequentially presented at the same spatial location (see Movie 2 and Fig. 3A).

**Figure 3.**
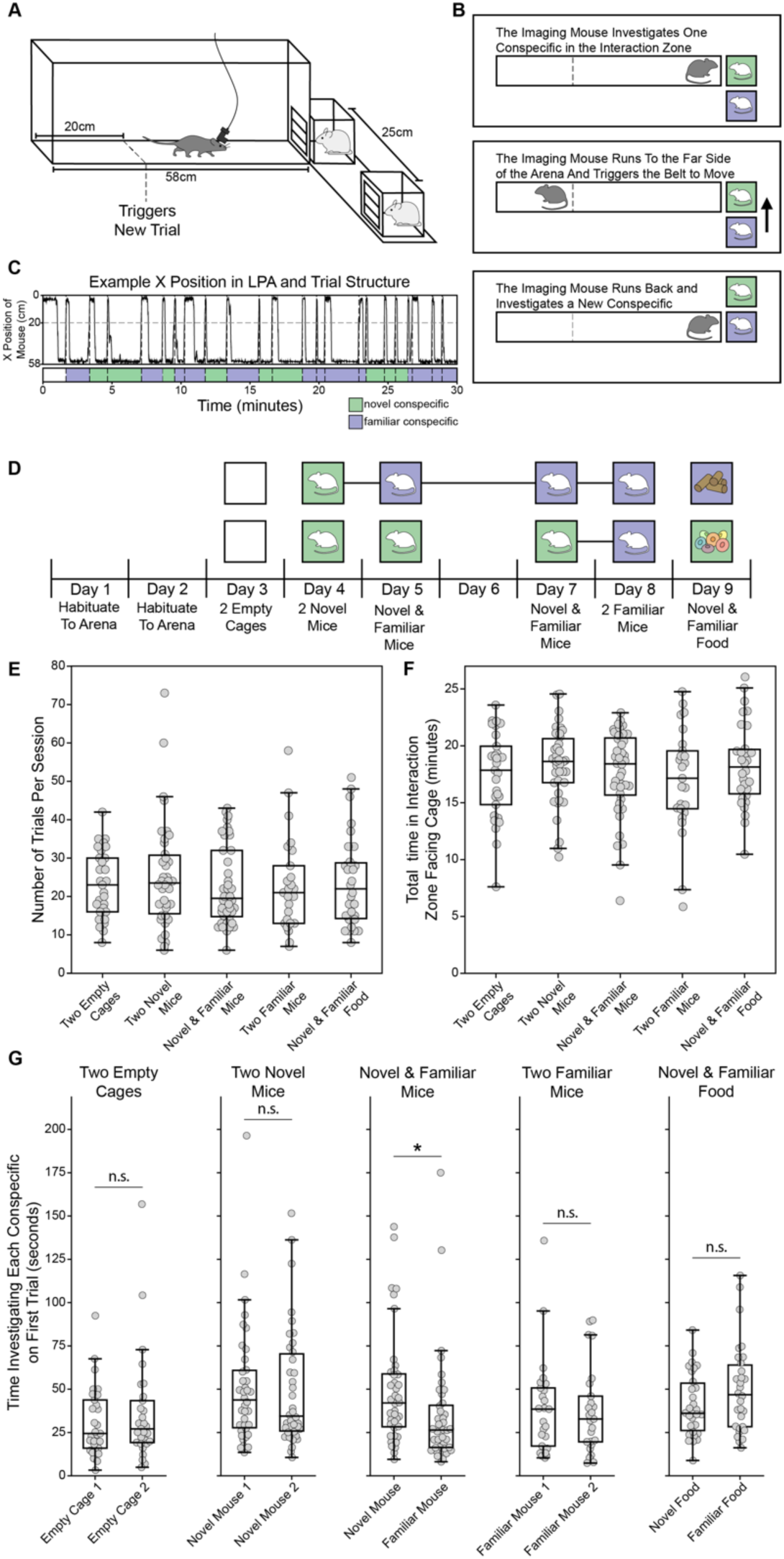
Linear Presentation Assay to Dissociate Social and Spatial Encoding. **A)** Schematic of the linear presentation assay (LPA). The imaging mouse (gray) can freely explore the linear arena. On the right end of the linear arena there is a barred window through which the imaging mouse can interact with a stimulus cage presented. An automated conveyor belt allows us to present multiple stimulus cages at the same spatial location. **B)** Schematic depicting the imaging mouse entering the trigger zone to initiate a new trial and trigger presentation of a new stimulus cage. **C)** Example time series of the imaging mouse’s position in the arena during a 30-min session on the LPA. Below, the corresponding time series of stimulus cage presented at the window is shown. The dashed line indicates instances when the imaging mouse triggers a new trial to begin. **D)** Timeline of behavioral recordings on the LPA. The colored boxes indicate the identity of the stimuli being presented for a given behavioral session. A horizontal black line connecting two colored boxes indicates that the same individual was reused as a social target across subsequent sessions. **E)** The total number of trials completed on the LPA in both iCA1 and iCA3 across behavioral conditions. **F)** Total time spent by the imaging mice in the interaction zone oriented toward the conspecific during 30-min sessions. **G)** The length of time that the imaging mouse investigates each stimulus cage on the first presentation of each cage (Wilcoxon signed-rank test: 2 empty cages: W = 176, n = 29, p = 0.370, effect size r = 0.191; 2 novel mice: W = 343, n = 38, p = 0.898, effect size r = 0.024; novel/familiar conspecifics: W = 235, n = 40, p = 0.0187, effect size r = 0.427; 2 familiar mice: W = 137, n = 25, p = 0.508, effect size r = 0.157; Novel/familiar food: W = 168, n = 30, p = 0.185, effect size r = 0.277)

We track the imaging mouse’s position in real time, and the belt automatically advances to the next cage when the mouse crosses to the side of the arena opposite of the window (Fig. 3B,C). Although the same stimulus cage can be presented multiple times in a row, an anti-biasing algorithm discourages long repeats of the same cage. This design enforces a pseudo-trial structure in which distinct social stimuli are presented repeatedly and unpredictably at a fixed location. By dissociating social identity from space, the LPA offers a powerful framework for probing hippocampal representations of social recognition independent of spatial confounds.

We ran several experimental conditions using the LPA (Fig. 3D). After habituating the imaging mice to the assay for two days, we first recorded neural activity while the mice investigated two empty stimulus cages. This condition served as a control to test whether mice could discriminate subtle manufacturing differences between the stimulus cages. Mice were then run through the assay in a series of different configurations, including presentations of: (i) two novel conspecifics, (ii) one novel and one familiar conspecific, (iii) two familiar conspecifics, and (iv) novel (Froot Loops cereal) and familiar food (rodent chow). In the food condition, mice could not access the food directly but could smell it through the interaction window.

Each LPA session lasted for 30 minutes, and the total number of trials per session varied (Fig. 3E). We excluded any session with fewer than four trials with each stimulus to ensure reliable statistical comparisons. On average, the imaging mice spent the majority of each session in the interaction zone, oriented towards the stimulus (Fig. 3F). Each session began with a fixed presentation of one stimulus cage followed by the other, after which stimulus cages were presented in a randomized order. Periods when the imaging mouse investigated each cage were hand-scored for bouts of active investigation, defined as periods of visible sniffing and/or whisking. In the novel/familiar conspecific condition, during the first two trials, the imaging mice preferentially investigated the novel conspecific longer than the familiar conspecific (Wilcoxon signed-rank test: W = 235, n = 40, p = 0.0187, effect size r = 0.427). The imaging mice did not display a significant behavioral preference in any other conditions (Fig. 3G; full statistics reported in figure legend).

We next determined that both the iCA1 and iCA3 neurons exhibit strong spatial encoding in the LPA (Suppl. Fig. 6A-D). Across all experimental conditions, the iCA1 consistently contained a higher proportion of place cells than the iCA3 (Suppl. Fig. 6E). At the population level, we assessed whether neural activity could decode the mouse’s spatial location. SVM classifiers decoded the mouse’s location from both iCA1 and iCA3 activity at well above chance levels in every condition. Notably, spatial decoding accuracy was significantly higher in the iCA1 than in the iCA3 across most conditions (Suppl. Fig. 6F, G).

### The iCA3 More Strongly Encodes Social Novelty than the iCA1

Next, we used an ROC analysis to identify neurons that preferentially fired during investigation of one conspecific over another (Fig. 4A,B). In the two empty cages condition, 5.2% of iCA1 neurons and 6.3% of iCA3 neurons displayed a significant preference, consistent with the expected false-positive rate given our statistical threshold (Fig. 4C). Similarly low proportions of neurons (7-9%; Fig. 4C) displayed a preference in the two-novel and two-familiar conspecific conditions in both regions. In contrast, in the novel-familiar conspecific condition (Day 5 in Fig. 3D), a significantly greater proportion of iCA3 neurons (16.4%; Fig. 4C) displayed a preference compared to iCA1 neurons (8.6%, proportion z-test; z = 5.70, p < 0.001). Lastly, in the novel-familiar food condition, a larger subset of neurons showed preferences in both regions (14.9% in iCA1 and 12.4% in iCA3; Fig. 4C). Distributions of social AUC values revealed a range of stimulus response strengths (Suppl. Fig. 7A, B).

**Figure 4.**
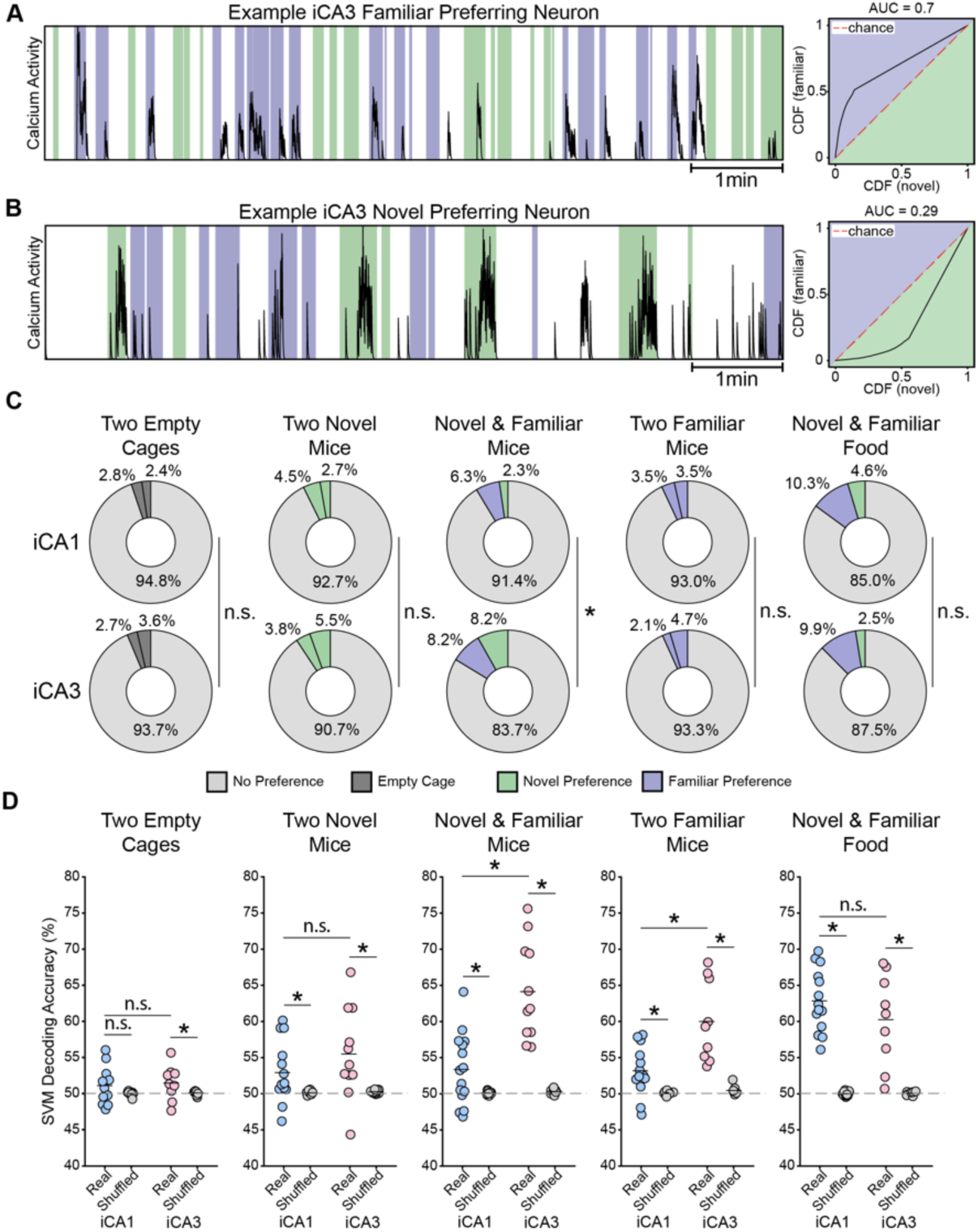
The iCA3 More Strongly Encodes Social Novelty Than the iCA1. **A)** Activity trace (left) and ROC (right) of an example iCA3 neuron that preferentially responds to the familiar conspecific while the imaging mouse investigates one novel and one familiar conspecific in the LPA. **B)** An example iCA3 neuron that preferentially responds to the novel conspecific. **C)** Proportion of neurons in each condition that display a preference for one stimulus cage in iCA1 mice (top) and iCA3 mice (bottom). Proportion z-tests were used to compare the fraction of significant neurons between regions (2 empty cages: z = 0.982, p = 0.326; 2 novel conspecifics: z = 1.779, p = 0.075; novel/familiar conspecifics (Day 5 in Fig. 3D): z = 5.704, p < 0.001; 2 familiar conspecifics: z = 0.208, p = 0.836; novel/familiar food: z = 1.6417, p = 0.101). **D)** SVM decoding accuracy in the iCA1 and iCA3 when decoding conspecific identity from population neural data. Each point represents the decoding accuracy of one mouse. Statistical significance between real and shuffled data, and between regions, was determined using hierarchical bootstrapping. In the two-empty cage condition, iCA1 decoding accuracy was not significantly above chance, whereas iCA3 decoding accuracy was (iCA1: Δ = 1.067%, p = 0.075, d = 1.951; iCA3: Δ = 1.375%, p = 0.034, d = 2.538). In the two-novel condition, both the iCA1 and iCA3 were significantly above chance (iCA1: Δ = 2.79%, p = 0.003, d = 3.630; iCA3: Δ = 5.21%, p = 0.002, d = 4.064). In the novel–familiar condition (Day 5 in Fig. 3D), both iCA1 and iCA3 decoding accuracies were significantly above chance (iCA1: Δ = 3.272%, p = 0.001, d = 3.733; iCA3: Δ = 13.566%, p < 0.001, d = 10.458). In the two-familiar condition, both iCA1 and iCA3 decoding accuracies were significantly above chance (iCA1: Δ = 3.14%, p < 0.001, d = 4.825; iCA3: Δ = 9.55%, p < 0.001, d = 7.524). In the novel– familiar food condition, both iCA1 and iCA3 decoding accuracies were significantly above chance (iCA1: Δ = 12.865%, p < 0.001, d = 16.281; iCA3: Δ = 10.19%, p < 0.001, d = 7.236). Decoding accuracy was significantly higher in the iCA3 than in the iCA1 only for the novel–familiar (mean difference = 10.508%, hierarchical bootstrap p < 0.001, Cohen’s d = 6.782) and two-familiar conditions (mean difference = 6.816%, hierarchical bootstrap p < 0.001, Cohen’s d = 4.832). Significant effects are indicated by asterisks; full test statistics and exact p-values are provided in Supplementary Data 1.

Next, we used binary SVMs to decode the stimulus targets from population activity in the iCA1 and iCA3. Statistical results are provided in the figure legend. In the two-empty-cages condition, neither the iCA1 nor the iCA3 achieved strong decoding accuracy, although mean iCA3 decoding accuracy was slightly above chance (Fig. 4D). This finding suggests that the empty cages were not robustly discriminated at the population level in either region.

In the social conditions (two-novel, novel/familiar, and two-familiar conspecifics), both the iCA1 and iCA3 displayed decoding accuracy significantly above chance. Decoding accuracy was significantly higher in the iCA3 than the iCA1 in the novel/familiar conspecific condition (Day 5 in Fig. 3D)(Δ = 10.51%, hierarchical bootstrap p < 0.001, Cohen’s d = 6.78) and the two-familiar conspecific condition (Δ = 6.82%, hierarchical bootstrap p < 0.001, Cohen’s d = 4.83) (Fig. 4D).

Decoding accuracy was also significantly above chance in the novel/familiar food condition in both the iCA1 (Δ = 12.85%, hierarchical bootstrap p < 0.001, Cohen’s d = 16.28) and iCA3 (Δ = 10.19%, hierarchical bootstrap p < 0.001, Cohen’s d = 7.24). This is particularly striking in the iCA1, where food decoding was significantly higher than decoding accuracy in all other conditions (Suppl. Fig. 7E).

We performed several controls to validate our decoding results. First, we replicated all decoding analyses using weighted F1 scores as the performance metric instead of accuracy and observed similar trends (Suppl. Fig. 7C). Second, we asked how decoding performance scaled with population size by training classifiers on increasing subsets of randomly sampled neurons within each mouse (Suppl. Fig. 7D). In iCA3, decoding of novel and familiar conspecifics was significantly above chance with as few as 10 neurons and improved as additional neurons were included. In contrast, iCA1 decoding hovered slightly above chance regardless of population size, consistent with our findings that social information is more strongly represented in the iCA3 relative to the iCA1.

Additionally, we allowed imaging mice to freely interact with conspecifics in an open arena to test whether the iCA1 and iCA3 similarly encode social recognition information during naturalistic dyadic interactions. Imaging mice investigated one novel and one familiar conspecific, one at a time, in alternating trials (Suppl. Fig. 8A). Imaging mice investigated the novel conspecific significantly more than the familiar conspecific (Suppl. Fig. 8B), consistent with prior findings in other assays (Fig. 1F; Fig. 3G). A significantly larger proportion of iCA3 displayed a social preference compared to iCA1 neurons (Suppl. Fig. 8C). At the population level, binary SVMs trained on iCA3 neural data had significantly higher decoding accuracy compared to the iCA1 (Suppl. Fig. 8D). Together, these results indicate that the regional differences observed in the SDT and LPA (Fig. 2H; Fig. 4D) are robust across behavioral assays and are also present when mice interact freely with conspecifics, rather than only when investigating them through a gate.

We also examined whether social and spatial decoding accuracy showed any topographical organization within the intermediate hippocampus (Suppl. Fig. 9). We overlaid each mouse’s decoding accuracy from the novel–familiar condition of the LPA onto the corresponding imaging plane reconstructions shown in Figure 1C. We did not observe any obvious topographical patterning of decoding accuracy beyond the overall regional differences between the iCA1 and iCA3 described above. Likewise, we found no differences in the decoding accuracies between straight and prism lens implants, although the sample size was not sufficient to make definitive conclusions.

### Longitudinal Tracking of Social and Spatial Representations in the Linear Presentation Assay

In the LPA, we observed functional distinctions between the iCA1 and the iCA3, with the iCA1 more robustly encoding spatial location and the iCA3 exhibiting stronger social representations. These differences were exaggerated when discriminating between novel and familiar conspecifics. We next asked whether either subfield carries a general social novelty signal. To test this, we tracked individual neurons across two novel/familiar LPA sessions separated by two days (Day 5 and Day 7 in Fig. 3D; Fig. 5A,B). The same familiar conspecific was presented in both sessions, whereas the novel conspecific was replaced with a new individual (Fig. 5C). A general novelty signal should therefore discriminate novel from familiar conspecifics in both sessions, even though the identity of the novel mouse differs. We also tracked spatial coding across the same sessions as a comparison.

**Figure 5.**
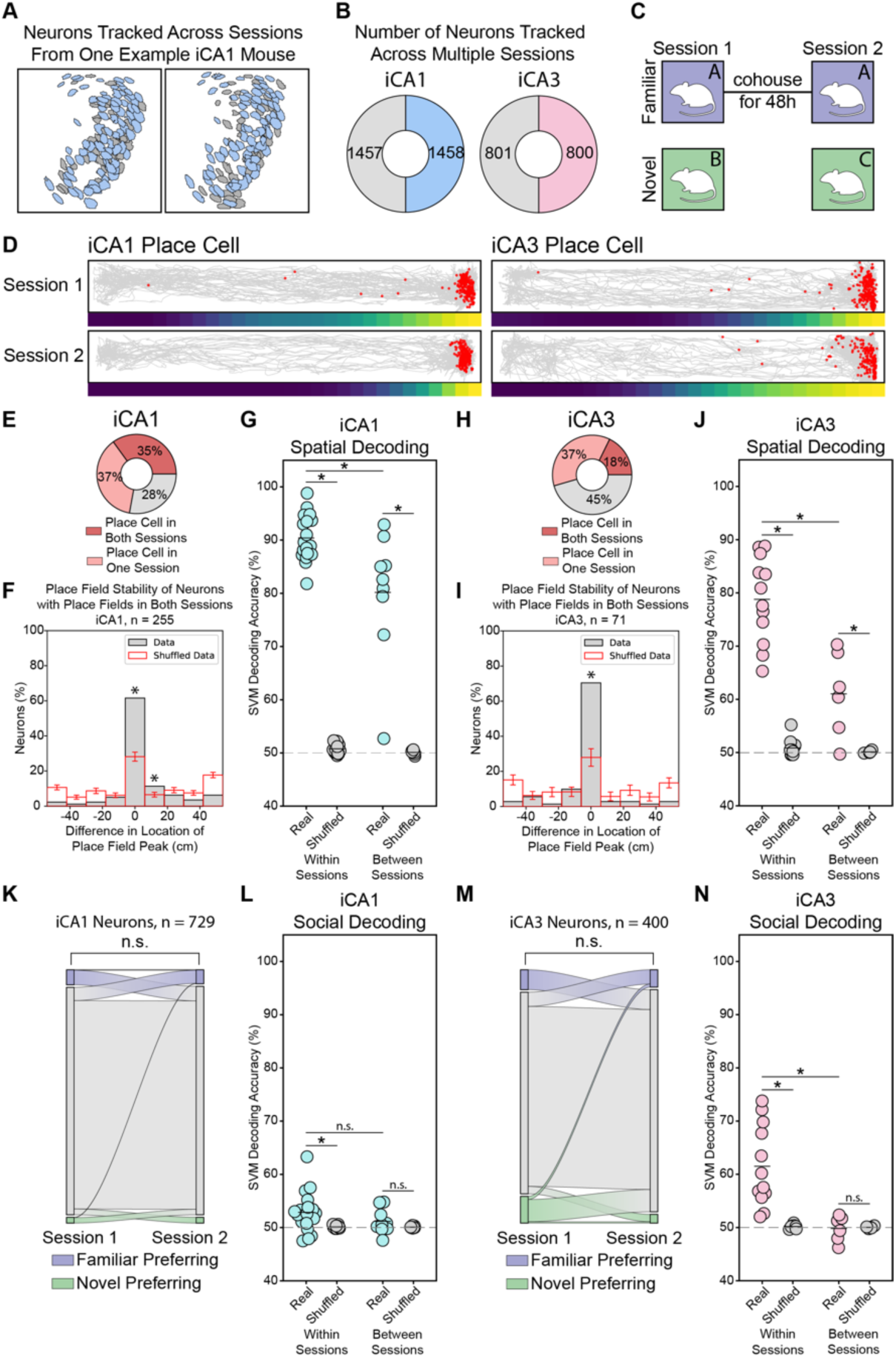
Longitudinal Tracking Dissociates Spatial and Social Coding in the Intermediate Hippocampus. **(A)** Example FOVs from the same iCA1 mouse across recording sessions on different days. Longitudinally tracked neurons are shown in blue; all other neurons are gray. **(B)** Number of neurons longitudinally tracked across sessions in the iCA1 (left) and iCA3 (right). **(C)** All subsequent longitudinal analyses use neurons tracked between one novel-familiar LPA session and a second novel-familiar LPA session using the same familiar conspecific paired with a new novel conspecific. **(D)** Two example place cells maintaining a stable place field across sessions. (**E**) Proportion of longitudinally tracked iCA1 neurons with a significant place field in each session. **(F)** Distance between place field peaks across sessions for neurons with significant place fields in both sessions. Significantly more iCA1 place cells mapped onto the same spatial location (gray) than expected by chance (red) (real = 61.57%, null = 28.22% ± 2.59%, p = 0.001, permutation test). **(G)** Cross-session decoding of spatial position in the LPA was restricted to longitudinally tracked neurons. Within-session decoders were trained and tested on data from the same session (independently for each of the two days); between-session decoders were trained on session 1 and tested on session 2. In the iCA1: within-session above chance (Δ = 39.60%, p < 0.001, d = 48.66); between-session above chance (Δ = 30.27%, p < 0.001, d = 11.93); within-session > between-session (Δ = 10.13%, p < 0.001, d = 3.78). **(H–J)** Same as (E–G), for iCA3. **(I)** Real = 70.42%, null = 27.96% ± 5.00%, p = 0.001. **(J)** Within-session above chance (Δ = 28.54%, p < 0.001, d = 18.09); between-session above chance (Δ = 11.56%, p < 0.001, d = 6.01); within-session > between-session (Δ = 17.51%, p < 0.001, d = 7.05). **(K)** Sankey plots depicting the proportion of neurons maintaining a social preference across the session pair in (C). In iCA1, observed stability (84%) did not significantly differ from the null distribution (mean = 84.73% ± 0.46%; permutation test, p = 0.95). **(L)** iCA1 social cross-session decoding. Within-session above chance (Δ = 2.72%, p < 0.001, d = 6.95); between-session at chance (Δ = 1.07%, p = 0.069, d = 2.00); within-vs. between-session not significant (Δ = 1.68%, p = 0.08, d = 2.56). **(M,N)** Same as (K,L), for iCA3. **(M)** Observed stability (73%) did not significantly differ from the null distribution (mean = 73.7% ± 0.96%; permutation test, p = 0.74). **(N)** Within-session above chance (Δ = 11.35%, p < 0.001, d = 9.98); between-session at chance (Δ = 0.20%, p = 0.56, d = 0.32); within-session significantly greater than between-session (Δ = 11.70%, p < 0.001, d = 9.03).

To confirm that neurons were tracked reliably, and that social- and spatial-selective neurons did not differ systematically in tracking quality, we compared registration confidence across social-selective, spatial-selective, and non-selective neurons. We found no significant difference between social-selective and non-selective cells (two-sample KS test: D = 0.127, p = 0.568) or between spatial-selective and non-selective cells (two-sample KS test: D = 0.052, p = 0.557), with all groups showing high median registration confidence (social = 0.939, spatial = 0.954) (Suppl. Fig. 10A, B).

First, we tested whether individual place cells within the LPA maintained consistent place field locations across conditions (Fig. 5D). We identified all neurons with significant place fields in both recording sessions (Fig. 5E,H), and then we calculated the Euclidean distance between the peaks of each field. In both the iCA1 and the iCA3, neurons maintained their spatial firing patterns more robustly than expected by chance (Fig. 5F,I).

To assess the stability of spatial representations at the population level, we trained SVMs to decode the mouse’s position in the LPA (i.e., opposite ends of the arena as seen in Supplementary Figure 6F) both within- and between-sessions. Within-session decoders were trained and tested on data from the same session (independently for each of the two days) whereas between-session decoders were trained on session 1 and tested on session 2. In both the iCA1 and iCA3, between-session decoders performed above chance levels (iCA1: Δ = 30.279, p < 0.001, d = 11.93, iCA3: Δ = 11.56, p < 0.001, d = 6.013). Between-session decoding accuracy was significantly lower than within-session decoding accuracy in both regions (iCA1: Δ = 10.125, p < 0.001, d = 3.78, iCA3: Δ = 17.509, p < 0.001, d = 7.05) (Fig. 5G,J).

To determine whether neurons maintained their social preference across sessions, we tracked neurons between the two conditions and compared the identity of the conspecific that they preferred. We classified each neuron as preferring the novel mouse, the familiar mouse, or showing no preference. We then quantified the proportion of neurons that retained the same preference across conditions. In both the iCA1 and the iCA3, neurons did not significantly maintain their social preference across conditions above chance levels (Fig. 5K,M).

To determine if there is a general social novelty representation at the population level, we replicated the between-session decoding approach outlined above. In the iCA1, within-session social decoding accuracy was significantly above chance, though only modestly (Δ = 2.721, p < 0.001, d = 6.949), while between-session decoding accuracy did not significantly differ from chance (Δ = 1.07, p = 0.069, d = 2.004) (Fig. 5L). In the iCA3, within-session decoding accuracy was significantly above chance (Δ = 11.35, p < 0.001, d = 9.984), while between-session decoding accuracy was not (Δ = 0.204, p = 0.564, d = 0.321), with a significant difference between within- and between-session decoding accuracy (Δ = 11.699, p < 0.001, d = 9.034) (Fig. 5N). Together, these findings suggest that while spatial coding in the iCA1 and iCA3 is stable both at the individual-neuron and population level, social novelty coding is comparatively more labile.

## Discussion

Social interactions require integrating information about who an animal encounters with where the encounter occurs, a computation thought to engage the intermediate hippocampus. Here, we provide the first recordings of endogenous activity in the iCA3 during social behavior and directly compare its function to the iCA1. Both iCA1 and iCA3 activity could discriminate between two novel conspecifics, but only modestly above chance, suggesting that both subfields carried some information about conspecific identity. However, iCA3 activity discriminated between novel and familiar conspecifics more clearly than the iCA1 and more clearly than it discriminated two novel conspecifics revealing a specific role in encoding social novelty. Decoders trained on one session did not generalize to a second session in which the novel conspecific was replaced, suggesting that the iCA3 supports real-time discrimination of a novel from familiar conspecifics rather than a general signal of social novelty. Together, these findings identify the iCA3 as a key site of social novelty encoding and establish a functional dissociation between the iCA1 and iCA3 in processing social recognition. These findings were made possible by a novel behavioral assay, the LPA, which controls for spatial position to isolate social representations.

### Extending the social recognition circuit beyond the CA2

The CA2 has been the hippocampal subfield most strongly implicated in social recognition, with lesion, inactivation, and recording experiments establishing the subfield as a critical hub for social computations^9–11^. The CA2 receives input from the CA3 and strongly projects to the CA1^9,32^. This connectivity positions CA2 as a potential conduit for transmitting social information through the hippocampal circuit. Our results reveal that social encoding extends beyond the CA2, engaging a broader network within the intermediate hippocampus.

To test how social recognition signals are embedded within broader hippocampal networks, we examined coding in the iCA1 and iCA3. In the SDT, we could decode spatial position more accurately than social recognition information in both the iCA1 and iCA3 (Suppl. Fig. 5A). Similar experiments recording from the CA2 have reliably decoded social recognition information, with spatial decoding accuracy varying across studies^8,11^. Furthermore, in our LPA paradigm, both iCA1 and iCA3 place cells mapped onto stable spatial locations across days (Fig. 5F,I), while CA2 place cells have been reported to rapidly remap across days^36^. These findings suggest a functional division of labor within the hippocampus. In the intermediate hippocampus, sparse social recognition signals are embedded within stable spatial frameworks, while in the CA2, stronger social representations are maintained in the context of more flexible spatial coding.

One potential mechanistic basis for this divergence lies in differences in synaptic plasticity across these regions. Both the CA1 and CA3 exhibit robust long-term potentiation^37–39^ whereas CA2 neurons are relatively resistant to conventional LTP induction^40,41^. Nonetheless, neuromodulatory inputs, such as oxytocin or vasopressin, can transiently enable plasticity in CA2 under certain conditions^42^. Reduced plasticity in the CA2 may favor the stability of abstract, identity-based social representations^43,44^, while greater plasticity in the CA1 and CA3 may support the integration of dynamic social information and spatially anchored representations.

### Conjunctive Encoding of Social and Spatial Features

In the SDT, a larger fraction of iCA1 and iCA3 neurons encoded conjunctive social–spatial information than social information alone (Fig. 2F,G). These findings underscore the intermediate hippocampus’s inherent bias towards contextual representations, where diverse forms of information are bound into relational codes^22^. In addition, population-level decoding revealed that joint social–spatial representations in the iCA1 and iCA3 were better represented than either component alone (Suppl. Fig. 5A), suggesting that the integration of these factors might be an active computational process rather than a random overlap between responses of individual neurons. This type of conjunctive coding might enable the hippocampus to link individuals to where they were encountered, facilitating the flexible retrieval of context-appropriate information.

Both the iCA1 and iCA3 project outside of the hippocampus^18,19,45^, raising the possibility that social–spatial codes are read and interpreted by downstream regions. Both subfields send strong projections to the lateral septum^20^, and we previously demonstrated that inhibiting ventral hippocampus–lateral septum neurons disrupts social novelty preference^46^. The iCA1 also projects to the medial prefrontal cortex (mPFC)^19^, and CA1–mPFC projections are likewise necessary for social novelty preference^47^. Notably, the mPFC itself contains neurons with conjunctive social–spatial selectivity^48^, similar to those observed in the iCA1 and iCA3, suggesting that the mPFC may inherit contextualized representations from the hippocampus.

### Limitations of the Linear Presentation Assay

To disentangle how the hippocampus represents social versus spatial information, we developed the LPA in which conspecifics are presented unpredictably and repeatedly at a fixed location. We show that mice are consistently motivated to investigate both social and non-social stimuli in this assay. Notably, we also found that both the iCA1 and iCA3 failed to robustly discriminate between two empty cages (Fig. 4D,E), suggesting that the presence of a meaningful stimulus is necessary to drive hippocampal representations. While the LPA provides a controlled environment for probing hippocampal encoding, it also has several limitations. Because the presentation of targets is externally controlled, experimental mice do not have the opportunity to choose what conspecific to approach and investigate. This limits the ability to assess whether a brain region encodes motivated social approach towards a specific individual. Nonetheless, the LPA’s ability to isolate social identity encoding independent of spatial encoding makes it a powerful complement to previous paradigms for disentangling social and spatial information.

### The iCA1 and iCA3 Subfields Differentially Encode Social Recognition Information

This study contributes to a growing body of literature that social recognition representations in the CA1 are topographically organized along the dorsal-ventral (DV) axis^25,49,50^. In rodents, inhibiting the dorsal CA1 (dCA1) does not disrupt social novelty preference in mice^12^, and endogenous recordings of the dCA1 have revealed less social modulation and social recognition encoding relative to more ventral CA1 neurons^12,51^. In contrast, recordings from the dCA1 in bats have revealed socially modulated place cells and neurons that encode both sex and identity^52–54^, suggesting that social representations may be more prominent in the dorsal hippocampus of some species. In this study, when controlling for spatial confounds, we find that few neurons in the intermediate CA1 display a social preference (Fig. 4C), and population-level decoders can decode conspecifics just above chance levels. This finding largely mirrors those of Okuyuma et al, 2016, who recorded from a similar location in the CA1 and also found that a small number of neurons display a social preference^12^. However, recordings from more ventral portions of the CA1 have revealed strong social modulation and strong sex, strain and identity encoding^51,55^. Together, these results suggest that there exists a functional gradient along the DV axis of the CA1, with the intermediate portion encoding sparse social recognition representations. Interestingly, we found that the iCA1 discriminated more robustly between novel and familiar food than between novel and familiar conspecifics. This result aligns with the well-established role of the CA1 in foraging and food-related memory^56–58^ and suggests that iCA1 may engage distinct coding strategies for social versus non-social forms of novelty. Future work directly comparing how hippocampal subfields represent novelty across social and non-social stimuli, including food, would help clarify whether these differences reflect stimulus-specific coding strategies.

Far less is known about how the CA3 contributes to social recognition. In the dorsal CA3 (dCA3), there are conflicting reports regarding its necessity for social novelty preference. Chiang et al., 2018 reported that inhibiting the dCA3 does not disrupt social novelty preference^13^ whereas Liu et al., 2022 found that optogenetic inhibition of dCA3-lateral septum neurons reduced social novelty preference^59^. In the intermediate/ventral CA3, Chiang et al., 2018 demonstrated that this region is necessary for social encoding but not for social recall. More recently, Li et al., 2025 expanded on this finding by showing that chemogenetic silencing basolateral amygdala inputs to intermediate/ventral CA3 also disrupts social novelty preference^60^. Building on these results, our study fills an important gap by providing the first recordings of endogenous iCA3 activity during social interactions. We provide evidence that the iCA3 contains robust and separable representations of novel and familiar conspecifics (Fig. 4D,E).

In this study, we found that both iCA1 and iCA3 share several functional properties. Each region contained conjunctive social–spatial representations (Fig. 2J), abundant place cells in both the SDT and LPA behavioral assays (Fig. 2F,G; Suppl. Fig. 6E), and spatial codes that were stable across sessions (Fig. 5F,G, I,J). Despite these similarities, the two regions diverged in how they represented social novelty. In the iCA1, social signals were sparse, and decoding performance was similar for discriminating between two novel conspecifics and between novel-familiar pairs. In contrast, the iCA3 showed significantly stronger separation for novel-familiar pairs than for two novel conspecifics (Suppl. Fig. 7E). One possible explanation for this difference relates to the underlying circuit architecture. The CA3 is often described as an autoassociative network, which is classically associated with pattern completion, the retrieval of a stored representation from partial or degraded input^61^. Such a mechanism could produce more categorical, attractor-like representations of familiarity than the more distributed coding observed in the iCA1. Consistent with this, we found that the iCA3 discriminated between two familiar cagemates with significantly higher decoding accuracy than the iCA1 (Fig. 4D). However, the interpretation of this finding is complicated by the fact that the two social targets were co-housed with the imaging mouse for an uneven length of time, raising the possibility that this result partly reflects residual novel/familiar discrimination. An interesting future direction would be to systematically explore how co-housing duration or behavioral dynamics between the imaging mouse and each conspecific affect the strength of identity-specific recognition representations.

### Social Novelty Representations in the Intermediate Hippocampus Remap Across Sessions

In both the iCA1 and iCA3, we found that a subset of neurons displayed significant place fields across two behavioral sessions (Fig. 5E,H), and those neurons exhibited highly stable spatial representations (Fig. 5F,I), consistent with longitudinal imaging recordings from the dCA1^62^. Such stability is expected from classical place cell studies^63^ and further indicates that our cross-session approach reliably captured meaningful neural dynamics. In contrast, social representations remapped across sessions (Fig. 5K,M). We suggest hippocampal social representations may support real-time social discrimination during the social encounter, rather than serving as a long-term storage site of social memory. This interpretation echoes the long-standing view that non-spatial hippocampal codes are more labile than spatial ones^22^.

One prominent counterexample to this interpretation is Okuyama et al. (2016), who reported that familiar-preferring neurons in the CA1 maintained a small but significant preference for familiar conspecifics over 24 hours. We find that after 48 hours, social representations remap almost completely, which may reflect a continuation of Okuyama’s findings with social novel/familiar representations continuing to drift over a longer timescale. Another notable exception to this pattern comes from recent work showing that ventral CA1 neurons can maintain highly stable representations of female odorants in male mice^64^. This discrepancy may reflect differences in social motivation, as our experiments used only same-sex conspecifics, which may be less behaviorally salient than opposite-sex conspecifics.

## Methods

### Experimental subjects

Male C57BL/6J mice (6–14 weeks old; Jackson Laboratory) were used for all iCA1 calcium imaging experiments (n=15). The iCA3 imaging cohort included two strains: a cohort of male C57BL/6J mice (6–14 weeks old; n=6 mice) in which imaging targeted the CA3 based on surgical coordinates refined during early experiments, and an additional cohort of male C57BL/6-Tg(Grik4-cre)G32-4Stl/J mice (6–14 weeks old; Jackson Laboratory; n=6 mice) used to more selectively target CA3 neurons. GRIN lens placements were verified post hoc via histological analysis in all implanted mice. To control for potential behavioral effects of lens implantation, a separate cohort of 28 C57BL/6J male mice (6–14 weeks old) received fluorophore (mCherry) injections into the intermediate hippocampus but were not implanted with lenses. These mice exhibited social preference comparable to implanted animals (Fig. 1F; Suppl. Fig. 2D), confirming that lens implantation did not alter social novelty preference. Age-matched male C57BL/6J albino mice carrying the white coat mutation (B6(Cg)-Tyrc-2J/J; Jackson Laboratory) were used as conspecifics in all experiments. All mice were housed under a reverse 12-hour light/dark cycle and all experiments were performed during the rodent night cycle. Experimental procedures were approved by the Emory University Institutional Animal Care and Use Committee.

### Viruses

The viruses used in this study were purchased from Addgene: AAV5-Syn-GCaMP6f-WPRE-SV40 (100835-AAV5; titer: 7×10¹² vg/mL), AAV5-Syn-Flex-GCaMP6f-WPRE-SV40 (100833-AAV5; titer: 7×10¹² vg/mL), and AAV5-CaMKIIa-mCherry (114469-AAV5; titer: 7×10¹² vg/mL).

### Calcium Imaging Stereotaxic Surgeries

6-8 week old mice (n = 15 iCA1 mice and 12 iCA3 mice) were anesthetized with 1-2% isoflurane and placed in a stereotaxic setup (Kopf). A microsyringe (Nanoject) was used to inject 500nL of the AAV5-Syn-GCaMP6f-WPRE-SV40 virus unilaterally into either the iCA1 or iCA3 region of the hippocampus of C57BL/6J mice at the rate of 2nL per second. For the Grik4-Cre mice, 700nL of AAV5-Syn-Flex-GCaMP6f-WPRE-SV40 (Addgene, injected titer of 1.3 × 1013 parts/ml) was injected into the iCA3 region. A week after viral injection, a 0.6 mm diameter, 7.3 mm length GRIN lens (Product ID: 1050-0004597, Inscopix) or a 1 mm diameter, 9.1 mm length prism GRIN lens with the lens face facing posteriorly (Product ID: 1050-004603, Inscopix) was implanted into the hippocampus. A metal cap was then cemented over the lens to protect it. Imaging mice were group housed following surgery. Three weeks following lens implant, a baseplate (Product ID: 1050-004638, Inscopix) attached to a miniature microscope (nVista, Inscopix) was positioned above the lens so neurons were in focus. The baseplate was then cemented in place (Metabond) and a custom-made baseplate cover was placed in the baseplate to protect the lens between imaging sessions

All stereotaxic coordinates are given relative to bregma. For iCA1, 0.6 × 7.3 mm straight lenses were implanted at ML ±3.7, AP –3.16, DV –3.5 mm, and 1 × 9.1 mm prism lenses at ML ±3.7, AP –2.9, DV –4.1 mm. For iCA3, 0.6 × 7.3 mm straight lenses were implanted at ML ±3.5, AP –2.8, DV –3.5 mm, and 1 × 9.1 mm prism lenses at ML ±2.8, AP –2.9, DV –4.3 mm. Viral injections were performed 0.1 mm ventral to the lens implant sites for straight lenses and 0.5 mm dorsal to the lens implant sites for prism lenses. For prism lens cohorts, bregma was measured relative to the center of the bottom edge of the prism face.

### Behavioral Assays

#### Social Discrimination Test for Novelty Preference

The social discrimination chamber (58.5 cm x 25.5 cm) contains two cylindrical cages (7.5 cm diameter) on either end of the assay. The experimental mice were habituated to the arena without any conspecifics present for 10 minutes each day, for two days prior to data collection. On the third day, the experimental mouse was placed in the assay with a novel age-matched same sex conspecific and a familiar age-matched same-sex conspecific who was cohoused with the experimental mouse for at least 24 hours preceding the test. The positions of the novel and familiar conspecifics (left vs right) were counterbalanced to eliminate side preferences. The experimental mouse could freely explore both conspecifics for 10 minutes while the behavior was tracked using an overhead camera. The position of the experimental mouse was tracked using a custom trained DLC network^65^.

To determine if the experimental mice displayed a novelty preference, we then calculated the percent of time that the centroid of the imaging mouse was located within a 12 cm radius around the cage. We than performed paired two-tailed t-tests to compare time spent around the novel and familiar conspecific. In addition, we calculated a discrimination score for each animal by subtracting the time spent near the novel conspecific by time spent near the familiar conspecific, divided by the total time spent near both animals, in line with previous studies^46^

#### Free Interaction Assay to Assess Novelty Preference

The behavioral assay was conducted in a featureless 29 cm diameter cylindrical arena. Experimental mice were habituated to the arena without any conspecifics present for 10 minutes per day over two consecutive days. On the third day, the experimental mouse was placed into the arena alone and allowed to freely explore for 2 minutes. After this baseline period, a novel or a familiar conspecific was introduced. The two mice were allowed to interact freely for 10 minutes while interactions were recorded with an overhead camera. Each mouse underwent two 10-minute sessions on the same day, one with a novel conspecific and one with a familiar conspecific. The order of conspecific presentation was counterbalanced across mice. Mouse positions and postures were tracked using a custom-trained DLC network. Identity swaps between the two mice were corrected using custom Python code and an autoencoder trained on hand-labeled frames was used to correct tracking errors.

Social investigation was quantified using the SimBA software suite^32^. To train the SimBA classifier, several behavioral recordings were manually annotated using BORIS^66^. We defined two behaviors: anogenital investigation and general social investigation. These behaviors were later combined into a single “social investigation” category for analysis. SimBA was trained on the DLC outputs and manual annotations, and its performance was validated by comparing its predictions to three withheld, hand-scored datasets (Suppl. Fig. 2B, C).

To determine whether experimental mice exhibited a novelty preference, we calculated the total duration of social investigation directed toward each conspecific. A paired two-tailed t-test was then performed comparing the percentage of time each mouse spent investigating the novel versus the familiar conspecific.

### Free Interaction Assay for Calcium Imaging Recording Sessions

Before imaging sessions, mice were habituated to the assay for two days, during which a tethered dummy miniscope was attached to the baseplate. The imaging mouse was placed into the arena for one minute, after which one familiar albino male mouse (cohoused for 24 hours) and one novel albino male mouse were presented in alternating three-minute trials separated by one-minute inter-trial intervals. Seven trials were completed in total (four with the familiar mouse and three with the novel mouse). Social targets were manually lowered into the arena at trial start and removed at trial end. The imaging mouse remained in the assay for the entire recording, and the arena was not cleaned between trials.

Instances of social investigation were annotated as described above. Wilcoxon signed-rank tests were used to determine whether imaging mice displayed a behavioral preference for one of the social targets (based on total investigation time across the first three trials with each target). For all imaging data analyses, including the single-neuron ROC and population-level SVM decoding, analyses were limited to frames in which the imaging mouse was actively investigating a social target. The single-neuron ROC and population-level SVM decoding analyses followed the same procedures as for the LPA (see *Linear Presentation Assay Receiver Operator Characteristics* and *Linear Presentation Assay Stimulus Decoders* below).

### Social Discrimination Tests for Calcium Imaging Recording Sessions

The social discrimination chamber (70cm x 70cm) contained two square stimulus cages (8cm x 8cm) positioned in opposite corners (top right and bottom left corners). Behavior was recorded under dim lighting conditions, with red-wavelength LED lights used to enhance video visibility while minimizing visual input for the mice. To provide visual spatial cues, high-contrast geometric shapes were affixed to one wall to serve as distal landmarks during navigation.

A tethered dummy miniscope attached to the baseplate was used to habituate the mouse (10 min x 2 days prior imaging). On the third day, imaging mice investigated two novel age-matched same-sex albino C57 mice on opposite corners of the assay. The imaging mice freely explored both conspecifics for 20 minutes and then we briefly removed the imaging mouse, cleaned both corners using ethanol, swapped the positions of the conspecifics and then returned the imaging mouse to freely explore the conspecifics for another 20 minutes. The swap generally lasted around 1 minute. On the fourth day, imaging mice investigated one novel albino C57 mouse and one familiar albino C57 mouse who was cohoused for 24 hours before data collection.

The imaging mouse had to interact with each conspecific a minimum of two trials both before and after the swap for sessions to be included in the analysis. To identify discrete trials, we drew a diagonal line between the two unoccupied corners of the arena, dividing the arena into two triangular regions. Each time the imaging mouse crossed this line and approached one of the conspecific cages was considered a new trial.

### The Linear Presentation Assay for Calcium Imaging Recording Sessions

The LPA consists of a long, narrow behavioral arena (57.5cm x 9cm) with a barred window on one end. Positioned perpendicular to the linear arena is a motorized conveyor belt (GVM GT-60D slider), which holds three small plexiglass cages (7.5cm x 7.5cm), referred to as stimulus cages. The stimulus cages are spaced evenly along the conveyor belt with 5cm gaps between each cage. The conveyer belt can move to align one of the stimulus cages with the window on the linear arena. When aligned, the stimulus cage and the linear arena window are separated by ∼1cm, allowing the imaging mouse and any mouse placed in the stimulus cage to make whisker contact and engage in olfactory investigation. The imaging mice are allowed to habituate to the linear arena while tethered using a dummy microscope (10min x 2 days). During experimental sessions, the imaging mouse was placed into the linear arena and was allowed to freely explore. Two stimuli are housed within the stimulus cages on the conveyor belt, while the center stimulus cage was always left empty.

We performed a series of social conditions using different combinations of conspecifics (age & sex-matched) such as two novel conspecifics, one novel conspecific and one familiar conspecific, and two familiar conspecifics. Familiar conspecifics were always cagemates who had been cohoused with the imaging mouse for at least 24 hours before data collection. In a separate experimental condition, we tested non-social novelty encoding by placing weight boats containing novel food (Froot Loops cereal) and familiar food (standard rodent chow). Lastly, in a control condition, we left all stimuli cages empty.

We recorded the behavior of the imaging mouse using an overhead camera, and we tracked the position of the imaging mouse in real time using a custom-written Bonsai script^67^. We defined the 10 cm area on the end of the linear chamber next to the window as the investigation zone. Furthermore, we defined the 20 cm zone on the opposite end of the linear chamber (the area furthest from the window) as the trigger zone (Fig. 3A). At the start of each recording session, the center stimulus cage (which is always empty) is aligned with the window in the investigation zone. When the imaging mouse moves from the investigation zone to the trigger zone, a custom pseudo random python function selects which stimulus cage to present at the window. Then custom-written bonsai code activates a Pulse Pal (Product ID: 1102, Sanworks) which sends a TTL pulse to trigger the conveyer belt to move, introducing a new stimulus cage to the window. The imaging mouse could then return to the investigation zone to investigate the new stimulus present.

We ran the LPA for 30 minutes. We defined a trial to be when the conveyer belt moves and the imaging mouse runs into the investigation zone to investigate the stimulus. The total number of trials and trial duration vary as function of the imaging mouse’s behavior for each recording session. After each session, behavioral videos were processed using a custom-trained DLC network to determine the position and orientation of the imaging mouse for further analysis.

To pseudo randomize stimulus cage presentation, we implemented a history-biased selection algorithm. While the same stimulus cage can be presented multiple times in a row, the algorithm reduced the likelihood of consecutive repeats and prevented any single stimulus cage from being presented more than four times consecutively. When repeats did occur, the conveyer belt would move to one side before quickly moving back to the cage that was just presented. This controls for mechanical noise emitted by the belt movement and prevents the imaging mouse from predicting which cage will be presented before entering the investigation zone. We excluded imaging sessions from data analysis if the imaging mouse failed to thoroughly investigate both stimulus cages. We required that the imaging mouse trigger a minimum of 4 trials with each stimulus. Additionally, we required that the imaging mouse had to be in the investigation zone and oriented towards the stimulus for a minimum of 5 seconds to be counted as a trial.

To determine if the imaging mice preferentially investigated one stimulus cage over the other, we hand-annotated instances of investigation using the Boris software^66^. Investigation was identifiable by the mouse’s snout reaching through the gate while actively whisking. We combined behavioral data from iCA1 and iCA3 cohorts with LPA sessions from a separate cohort of imaging mice, as the behavioral protocol was identical across cohorts. In Fig. 3G, we limited our comparison of investigation times to the first two trials of each session because these are the only trials that have a fixed order; one cage is randomly presented first and then the other is presented. In all subsequent trials, the same cage may be presented multiple times in a row, and repeated exposure to the same conspecific could itself reduce investigation independent of its novel/familiar status. Restricting the analysis to the first two trials avoids this confound and also ensures that the novel conspecific remains novel at the time of testing.

### Endoscopic calcium imaging data collection and analysis

During imaging sessions in both the SDT and the LPA, a miniscope (nVista, Inscopix) was placed on top of the baseplate of each mouse and fastened in place using a small screw. The LED power on the miniscope was set to 0.5 and the analog gain on the image sensor was set between 1.4 and 1.8. Calcium imaging data was recorded at 20Hz using the Inscopix data acquisition software. After data collection, imaging recordings were spatially downsampled by a factor of 4 and motion corrected using Inscopix software. The CNMFe algorithm^68^ was then used to identify individual neurons and extract their florescence traces. Florescence traces for each animal were then aligned to behavioral events. Custom Python software was used for all imaging data analysis.

### Histology and Microscopy

After all experiments were complete, mice were euthanized and perfused with 0.5% PBS followed by 4% paraformaldehyde (PFA). The brains were extracted and fixed in 4% PFA overnight at 4° F before being transferred to a 30% sucrose solution for another 24 hours. Brains were sliced at 50 µm using an Epredia HM430 microtome and the slices were mounted in DAPI solution (Product ID: 0100-20, SouthernBiotech). The brains of imaging mice that were implanted with a prism lens were sliced sagittally to better determine lens placement. All other brains were sliced coronally. Slices were imaged using florescence microscopy (Keyence BZ-X800).

### Place Cell Quantification in the Linear Presentation Assay

To identify place in the LPA, we first isolated time periods during which the imaging mouse was moving. In line with prior studies, we included frames where the moving average speed (computed in a 0.5-second window) of the mouse’s centroid exceeded 4 cm/s^28^. The LPA behavioral arena was divided into 10 spatial bins × 9 cm). The identity of the stimulus at the interaction zone was ignored when computing spatial information. Neural activity was preprocessed by smoothing the deconvolved calcium signal with a Gaussian kernel (σ = 75 ms). Next, each deconvolved trace was time binned by averaging the values in 200 millisecond bins. For each spatial bin in the behavioral arena, we calculated the total time that the mouse spent time in that bin and summed the total neural activity when the imaging mouse occupied each bin. We excluded any spatial bins that the mouse occupied for less than 1 second. We then computed the average neural activity per bin and normalized this by the occupancy to create spatial firing maps.

We calculated the spatial information content of each neuron using the following equation^69^:

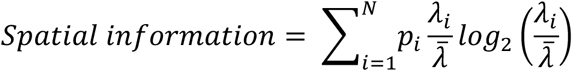

where *p_i_* is the occupancy probability of spatial bin *i*, *λ_i_* is the mean neural activity in that bin, and 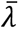 is the mean activity across all bins. To determine whether spatial information exceeded chance for each neuron, we generated a null distribution by circularly shifting each neuron’s activity trace 1000 times, using a minimum shift of 10 seconds. Neurons whose spatial information exceeded the 95th percentile of the null distribution were classified as place cells. To calculate the location of place fields, we identified all spatial bins whose values exceeded 50% of the global maximum activity in line with previous papers^28^. Spatial maps were smoothed for visualization only (Suppl. Fig. 6A-D); spatial information was computed on the unsmoothed data. In the LPA, we defined the place field peak as the spatial bin with the highest activity value.

### Social Discrimination Test Receiver Operator Characteristics

To determine whether individual neurons preferentially fired for one conspecific over another in the social discrimination test we used a receiver operating characteristic (ROC) analysis. We used the same approach to determine if neurons preferentially fired for the space where the two conspecific cages were present.

We first identified frames when the imaging mouse investigated each conspecific using the position information of the imaging mouse tracked using DLC^65^. Frames were included in data analysis if the mouse’s centroid was located within an 11 cm radius of one of the corners containing a social stimulus cage. To ensure that the mouse was oriented toward the stimulus, we tracked the nose and ears of the mouse and computed a head direction vector from the midpoint between the ears to the nose. Frames were only included if the stimulus cage lay within a 240° cone centered on this head direction vector. We then divided all valid frames into discrete trials as described above.

We grouped the trials into four different conditions, when the imaging mouse investigated each conspecific before and after the swap. To prevent confounds between social identity and spatial location in the ROC analysis, we equalized the number of frames across all four social–spatial conditions using a round-robin sampling approach. Frames from longer trials were truncated to ensure that each condition contained an equal number of frames. This trial matching ensured that neither conspecific nor spatial location was overrepresented in the binary ROC comparisons.

To assess whether individual neurons discriminated between conspecifics or spatial locations, we re-grouped the four conditions into two binary classifications. For social ROC, data from both spatial locations were combined for each conspecific, yielding two social conditions. For spatial ROC, data from both conspecifics were combined by location, yielding two spatial conditions. For each neuron, we then extracted the normalized fluorescence trace during all frames of investigation and computed the area under the ROC curve (AUC) using the roc_auc_score function from the sklearn.metrics package in Python^70^. The comparison direction (i.e. the condition labelled as the positive class) was held constant across all neurons and as a result, AUC values below 0.5 indicate a neuron preferentially fires to the alternative condition in line with prior literature^71,72^. To assess significance, we circularly shifted each neuron’s activity 1000 times to generate a null distribution of AUC values. Each shift was randomly drawn from a uniform distribution spanning 10-90% of the length of the session to prevent small shifts from wraparound. We considered a neuron to show a significant preference if its real AUC fell outside ±2 standard deviations of the null distribution.

We classified neurons as social–spatial in the following way. For each neuron, we computed the AUC–ROC value between each conspecific in the top right corner and AUC-ROC value between each conspecific in the bottom left corner. We then calculated the difference in AUC values between these locations. This “difference-of-differences” interaction value was compared to a null distribution generated by circularly shifting each neuron’s activity trace. Neurons whose observed interaction exceeded two standard deviations above the null distribution were classified as social–spatial neurons. As a post hoc confirmation, we also required that these neurons showed significantly greater activity in one social–spatial condition compared to all others (permutation test, p < 0.05). Additionally, any neurons that exhibited both a significant social main effect and a significant spatial main effect were also classified as social–spatial neurons.

### Linear Presentation Assay Receiver Operator Characteristics

To determine whether individual neurons preferentially fired for one stimulus over another in the LPA, we applied the same ROC-based analysis described above, with a few key differences in data preparation.

First, we tracked the imaging mouse using DLC to index frames when the centroid of the imaging mouse was in the predefined interaction zone. We also indexed frames when the imaging mouse was oriented toward the stimulus, determined by the vector from the centroid to the nose of the animal. We then compiled two lists of frames: one for interactions with each stimulus. We then trial matched the lists by deleting excess frames from the end of the longer list. The normalized fluorescence trace for each neuron was then extracted from these timepoints, and ROC AUC values were computed using the same approach as described above.

### Social Discrimination Test Support Vector Machines for Social and Spatial Decoding

We used support vector machines (SVMs) to decode conspecific identity from population neural activity. The entire imaging session was divided into one-second time bins. We identified time bins during which the imaging mouse was both near one of the conspecifics and oriented toward the conspecific as described above. Only time bins in which all frames met both spatial and orientation criteria were included in decoding. For each neuron, we extracted the average activity within each valid time bin and z-scored the resulting values across all valid time bins.

To prevent information leakage between training and testing data, we grouped valid time bins into discrete trials, as described above. Each trial was labeled as one of four classes: Conspecific A before swap, Conspecific B before swap, Conspecific A after swap, or Conspecific B after swap. We performed 100 iterations of the following procedure: First, we randomly shuffled the trial order within each class. Next, the number of time bins per class was trial-matched by truncating excess time bins to ensure equal representation across all classes. Each class was divided into five equal folds; four folds were used for training and one for testing.

When decoding social identity, time bins were pooled across spatial locations for each conspecific, yielding two binary classes: Mouse A and Mouse B. When decoding spatial location, time bins were pooled across conspecifics, resulting in two classes corresponding to the top right and bottom left corners of the arena.

We performed nested 3-fold cross-validation within the training set of each iteration to tune regularization parameters. We performed a grid search over a range of L2 regularization strengths. SVM classification was performed using a linear kernel with the LinearSVC function from the sklearn package in Python^70^. For each mouse, we report the average test decoding accuracy across all folds and all iterations. A pipeline of this decoding procedure is shown in Supplementary Figure 4.

To determine whether decoding accuracy significantly differed from chance, we generated a null distribution of decoding accuracies on chunk-shuffled calcium imaging data. We first binned the calcium imaging traces into 2-second time bins and then we shuffled the bins for each neuron independently of all other neurons. This shuffling approach preserved the structure of the calcium events while disrupting the temporal relationship to behavioral events. For each mouse we then ran 100 iterations of the support vector machine decoder as described above and we reshuffled the calcium imaging data on every iteration. We reported the average decoding accuracy across all iterations and all folds.

### Social Discrimination Test Social–Spatial Decoders

To decode all four social–spatial conditions simultaneously, we formatted the data as described above, with time bins trial-matched across the four experimental classes. Unlike the binary decoding approach used to classify either social identity or spatial location, this four-way decoder treated each unique social–spatial condition as a distinct class (Suppl. Fig. 4). We trained a multiclass support vector machine (SVM) classifier using a one-vs-rest strategy, as implemented in LinearSVC from the sklearn package in Python^70^. Performance was quantified using overall classification accuracy, computed as the proportion of correctly classified time bins across all four classes.

### Normalizing Decoding Accuracy in the Social Discrimination Test

To enable comparisons across decoding models (social, spatial, and social–spatial), decoding accuracy was normalized relative to chance performance. For each session, the decoding accuracy from each iteration (*DecodingAccuracy_true_*) was first centered by subtracting the mean decoding accuracy obtained from shuffled neural activity (*DecodingAccuracy_s_*_ℎ*uffled*_) and then scaled by the maximum possible improvement above chance:

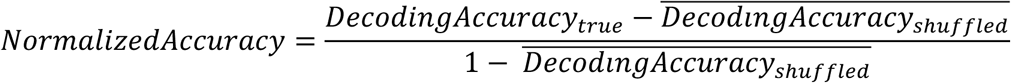

This normalization controls for differences in chance levels across sessions and models and rescales decoding performance relative to the maximum attainable value above chance, enabling direct comparison across models.

### Hierarchical Bootstrapping

To determine whether the decoding accuracies of multiple mice within an experimental condition significantly differed from chance, we implemented a hierarchical bootstrapping approach as described in Saravanan et al., 2020^73^. For example, we applied this method to test whether social decoding accuracy in iCA1 mice significantly exceeded that of shuffled controls. We performed 1,000 iterations of hierarchical resampling: on each iteration, we sampled mice with replacement, and for each sampled mouse, we resampled decoding iterations with replacement. We then computed the mean decoding accuracy across mice for each resampled dataset, resulting in a distribution of 1,000 bootstrapped mean accuracies.

To evaluate statistical significance between real and shuffled decoding distributions, we calculated one-sided p-values using a joint probability matrix approach. For comparisons between two real conditions (e.g., iCA1 vs. iCA3), we used two-sided p-values computed with the same method.

### Social Discrimination Test Decoding Accuracy Before and After the Conspecific Swap

To examine whether decoding accuracy varied systematically across the recording session, we analyzed decoder performance separately for time bins before and after the conspecific swap. We ran 100 iterations of the support vector machine decoder as described above, and for each time bin in the test set, we computed the average signed distance from the decision boundary. We then determined whether each time bin was correctly classified on average across all iterations and calculated the percentage of correctly classified time bins separately for the pre- and post-swap periods. We were particularly interested in whether overall above-chance accuracy masked asymmetric confidence.

### Linear Presentation Assay Stimulus Decoders

We implemented a similar decoding approach on the LPA to the approach outlined in the SDT. To start, we divided into the entire recording session into one second time bins. We identified time bins in which the centroid of the imaging mouse was in the interaction zone as previously described. We then formed a vector between the centroid of the imaging mouse to the nose (180° cone) to determine if it was oriented towards the stimulus. We then generated two lists of time bins that met these criteria, one for time bins when stimulus A was present and one when stimulus B was present. To trial-match time bins across stimuli, we randomly shuffled the trial order and truncated excess time bins from the condition with more data. Then we repeated the same binary support vector machine approached as described above. We used two approaches to determine if stimulus decoding accuracy was significantly above chance on the LPA. We used permutation tests to determine if the decoding accuracy of any individual mouse was significantly above chance. We considered a session to significantly decode the stimuli if the real decoding accuracy was above 2 standard deviations of the distribution of shuffled decoding accuracies. We utilized the hierarchical bootstrapping approach as described above to identify significant differences in population decoding across regions.

As an alternative performance metric to decoding accuracy, we also calculated weighted F1 scores, which account for both precision and recall while weighting each condition according to its frequency in the test set. To compute weighted F1 scores, we largely followed the decoding pipeline described above, with a few key differences. Most importantly, we did not trial-match the data, allowing us to include all available time bins from each condition. Weighted F1 scores were calculated using the f1_score function from the sklearn.metrics package in Python^70^, with the parameter average=’weighted’ to ensure that each class contributed proportionally based on its prevalence.

### Linear Presentation Assay Spatial Decoders

We used a SVM machine to decode the location of the imaging mouse in the LPA from population neural data. We divided the recording session into one-second time bins and identified bins in which the mouse’s centroid was within 14.4 cm of either the stimulus window or the opposite wall, corresponding to the outermost 25% ends of the arena. To ensure sufficient data for decoding, we required that the imaging mouse occupy each side of the arena over 4 discrete trials with a minimum of 5 time bins per trial. To trial match time bins for each location, for each decoding iteration, we randomly shuffled the order of the trials and then we deleted time bins from the end of the location with more data. Then we repeated the same binary support vector machine approached as described above. Lastly, we generated a null distribution by chunk shuffling calcium imaging data as described above and we assessed significance using the hierarchical bootstrapping approach as described above.

### Longitudinal Tracking Neurons Across Imaging Sessions

Neurons were longitudinally tracked across imaging sessions using a widely used^74–76^ probabilistic cell-tracking algorithm, CellReg package in MATLAB^77^. CellReg estimates the probability that a candidate pair of cells represents the same neuron (P(same)) based on spatial footprint correlation and centroid distance between the candidate neurons. To assess whether registration confidence differed systematically across cell types, we extracted the P(same) value for each pair of tracked neurons (Suppl. Fig. 10). All neuron matches were additionally manually inspected to confirm consistency in spatial footprint and centroid location. Neurons with ambiguous matches or poor-quality spatial footprints were excluded from analysis.

### Linear Presentation Assay Between-Session Social and Spatial Decoding

To decode either conspecific identity or spatial location across two imaging sessions, we used a SVM framework similar to the one described above. First, we identified all neurons that were successfully tracked across both sessions. If fewer than 15 neurons were tracked between a pair of sessions, the session pair was excluded from cross-session decoding.

We trained an SVM on the first session using only the subset of neurons tracked across both sessions. We repeated this training procedure 100 times with randomized trial splits, and for each neuron, we averaged its learned decoding weight and intercept across all iterations. These decoding weights and intercepts were then transferred to the second session. We projected the standardized neural activity from the second session onto the trained hyperplane using the weights and intercepts learned from the first session. We repeated 100 iterations of the SVM on the second imaging session using randomized trial splits, and we calculated the decoding accuracy on each iteration.

To assess significance, we generated a null distribution by performing 100 iterations of chunk shuffling on the test session only. On each iteration, calcium imaging traces from the second session were divided into 2-second bins, and the order of these bins was independently shuffled for each neuron. The decoder, trained on the unshuffled first session, was then applied to the shuffled test session. To assess whether decoding performance across mice within a given condition (e.g., iCA1 spatial decoding) was significantly above chance, we applied the hierarchical bootstrapping procedure described above. To evaluate the stability of representations across sessions, we also compared between-session decoding accuracy to a within-session control, generated exclusively using the subset of neurons that were tracked across session.

### Cross-Session Place Cell Analysis

To test whether place cells remap across sessions in the LPA, we first identified all neurons that exhibited significant place fields in two separate sessions. As described above, we divided the LPA arena into 10 discrete spatial bins (5.7 cm × 9 cm). For each neuron, we identified the bin with the peak activity in each session and we calculated the Euclidean distance between these peak spatial bins.

To determine whether the observed remapping exceeded what would be expected by chance, we randomly permuted neuron identities across sessions and recalculated the distance between peak bins for each shuffled pairing. This procedure was repeated 1,000 times to generate a null distribution of remapping distances.

To compare the real and shuffled distributions, we overlaid their histograms and assessed whether the real distribution was more strongly peaked around zero than chance would predict. Specifically, for each discrete distance value, we compared the percentage of real neurons at that distance to the null distribution and considered the difference statistically significant if it exceeded two standard deviations above the null-distribution mean for that bin.

### Cross-Session Social Cell Analysis

To assess whether neurons maintained their social preference across sessions on the LPA, we first identified all neurons that could be tracked across both sessions. We then compared each neuron’s preference (as determined by ROC analysis) in the two different sessions. Neurons were assigned to one of three categories in each session: preference for Mouse A, no preference, or preference for Mouse B. We calculated the proportion of neurons that retained the same preference across sessions. To evaluate statistical significance, we performed 10,000 permutations in which the preference labels from the second session were randomly shuffled across neurons. For each permutation, we recalculated the proportion of neurons with matching preferences, yielding a null distribution. We computed a one-sided p-value as the proportion of shuffled iterations in which the proportion of matching preferences equaled or exceeded the observed value.

### Statistical analyses

All statistical tests were performed using custom python scripts. All of the statistical tests that were performed and their corresponding values can be found in Supplementary Data 1.

## Supplementary materials

**Supplementary Figure 1.**
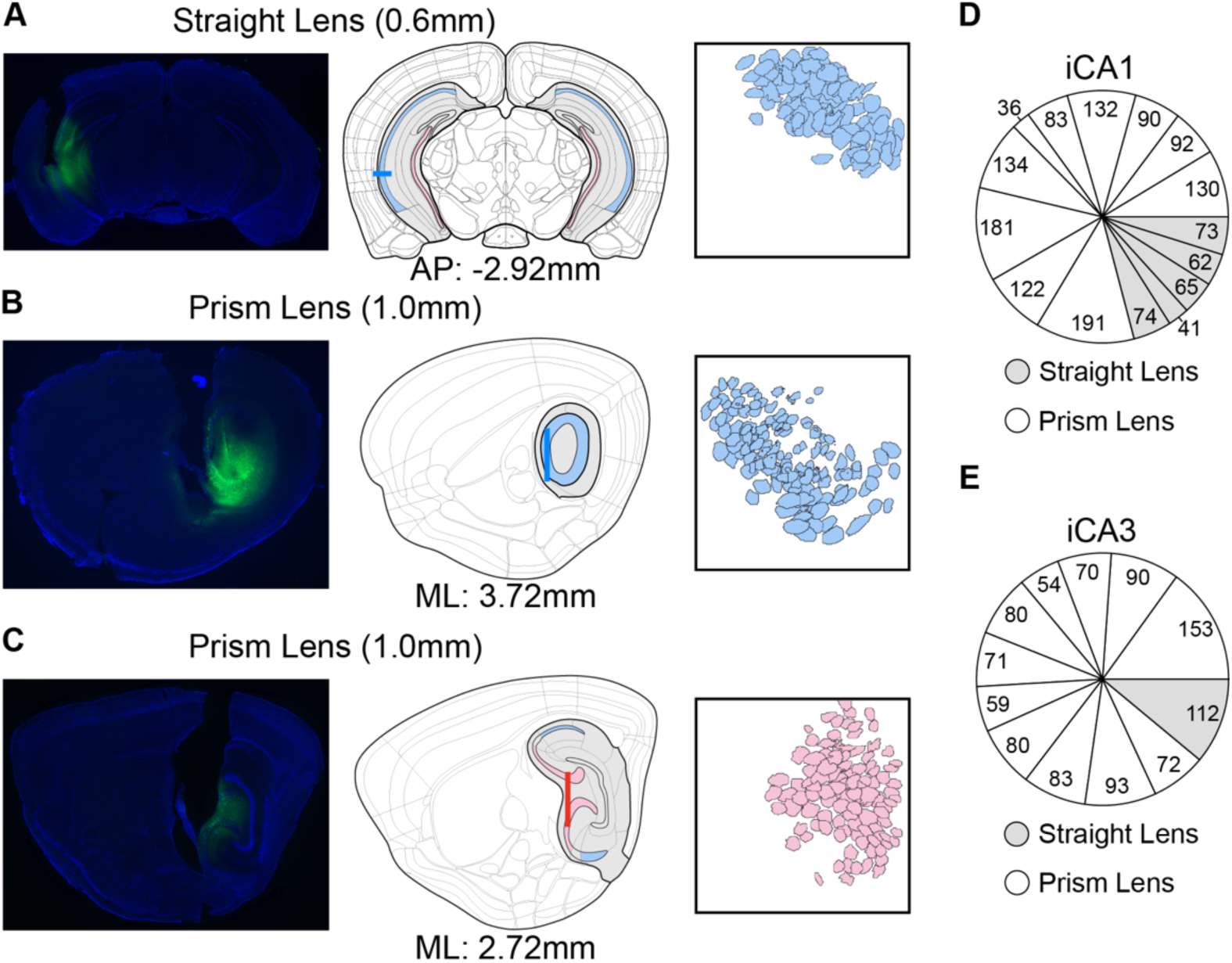
Imaging Histology. **A)** (left) Coronal section from a brain implanted with a 0.6-mm straight lens targeting the iCA1, showing GCaMP expression. (center) Coronal section at the corresponding imaging plane. (right) FOV from the same iCA1 imaging mouse. **B)** (left) Sagittal section from a brain implanted with a 1.0-mm prism lens targeting the iCA1, showing GCaMP expression. (center) Sagittal section at the corresponding imaging plane. (right) FOV from the same iCA1 imaging mouse. **C)** Same as (B), but targeting iCA3. **D)** Pie chart showing the distribution of recorded neurons across iCA1 imaging mice, separated by lens type. **E)** Same as (D) but for iCA3.

**Supplementary Figure 2.**
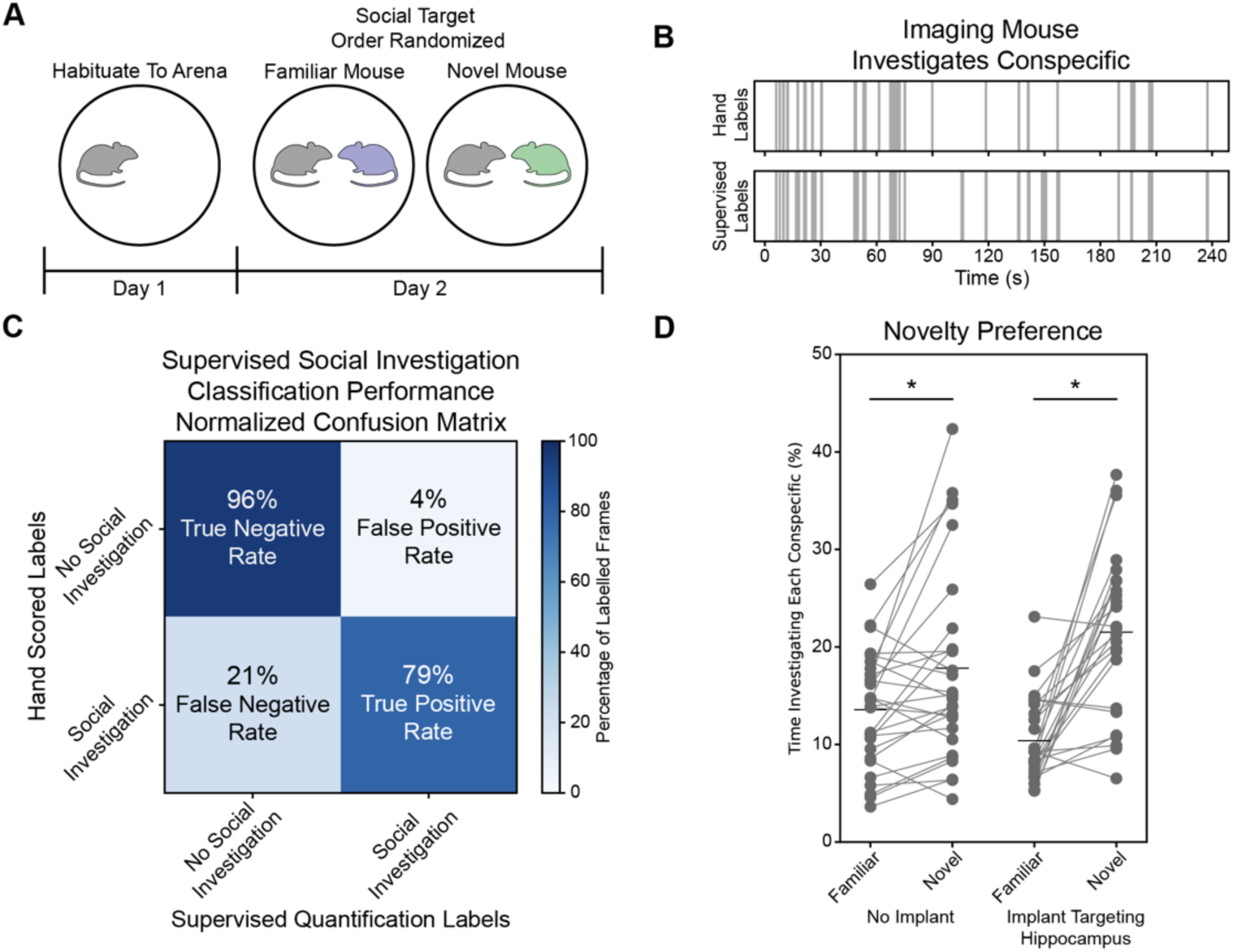
Imaging Mice Display a Novelty Preference in a Free Interaction Test. **A)** Timeline of the free interaction test. **B)** Comparison of hand-labeled social investigation (top) and SimBA-labeled social investigation (bottom). **C**) Confusion matrix showing SimBA performance on three hand-labeled behavioral recordings not included in the training set. **D)**) Mice preferentially investigated novel conspecifics over familiar conspecifics in lens implanted mice and a no lens control cohort (lens implanted mice: paired-test (t(23) = 5.776, p < 0.001, n = 25; no lens: paired t-test t(27) = 3.025, p = 0.005, n = 28).

**Supplementary Figure 3.**
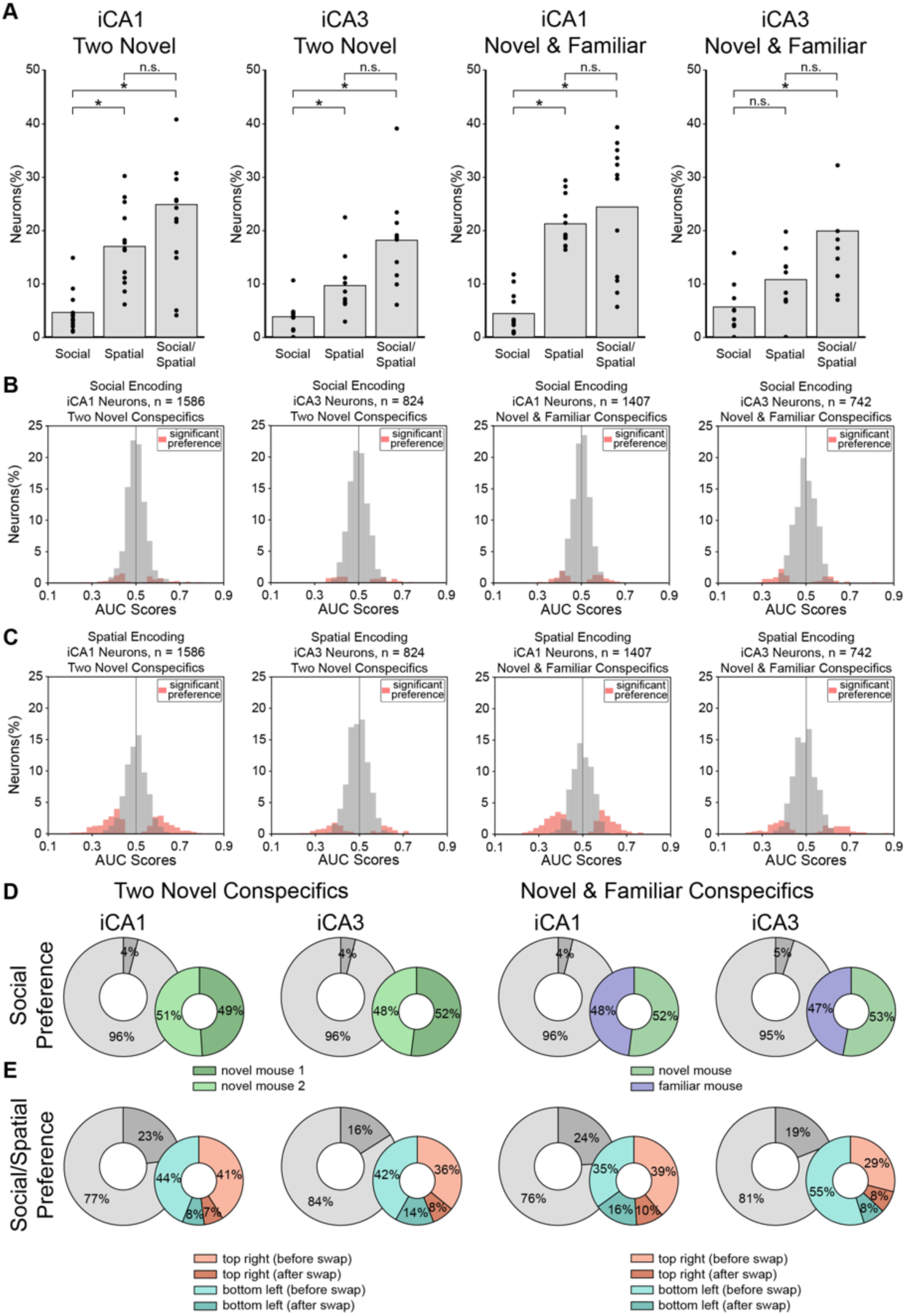
SDT Encoding Responses. **A)** Proportion of neurons showing a preference in the SDT plotted separately by mouse. Although individual datasets have lower statistical power, the trends are consistent with the population-level effects shown in Figure 2; full test statistics and exact p-values are provided in Supplementary Data 1. **B)** Distribution of social AUC values across all neurons in the SDT. **C)** Same as (B) but for spatial AUC values. **D)** Among neurons with a social preference, the proportion preferring each social target. **E)** Same as (D), but for neurons displaying a social–spatial interaction.

**Supplementary Figure 4.**
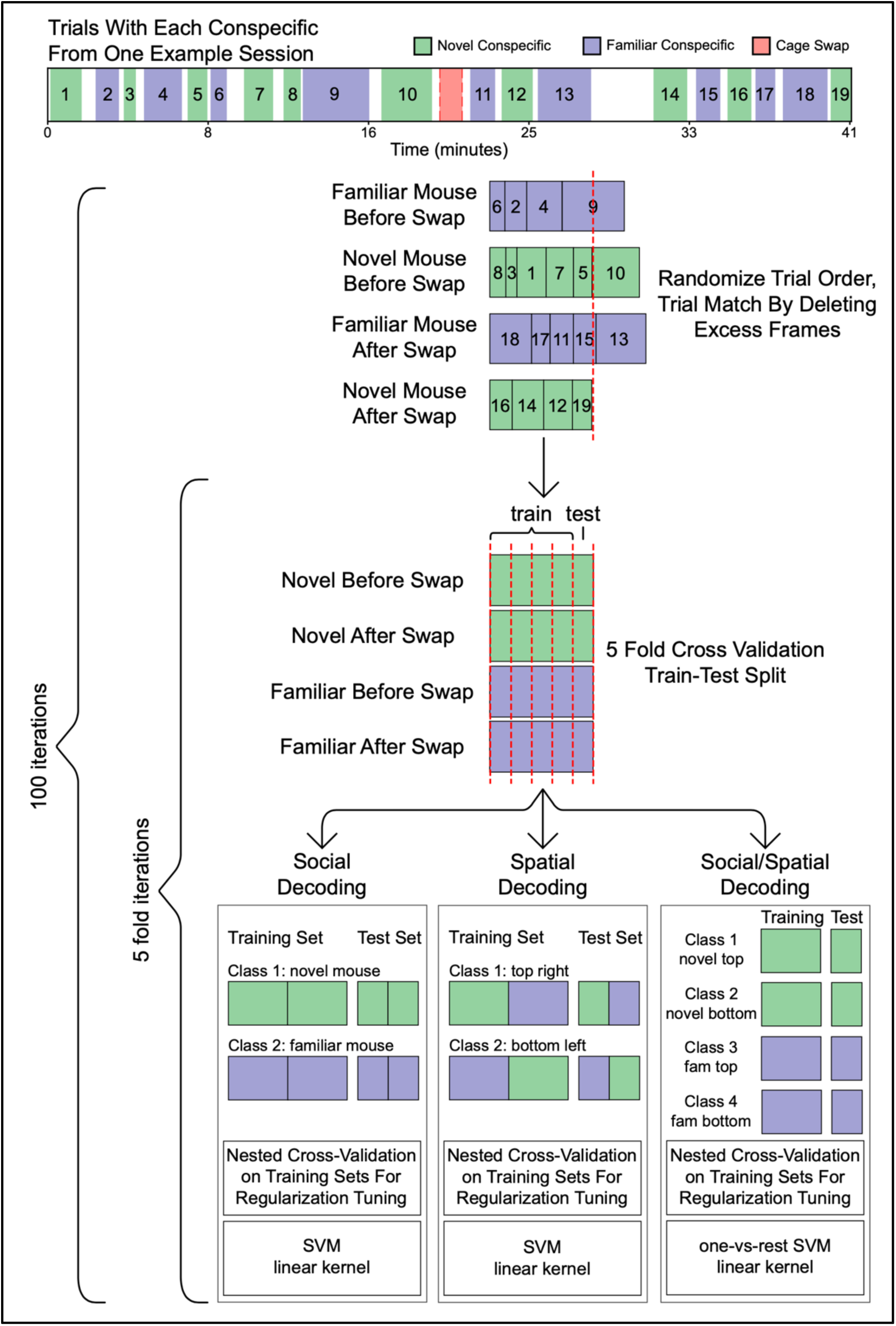
SDT SVM Decoding Approach. Workflow of the SDT decoding approach. Schematic illustrating the trial matching procedure and subsequent partitioning of the data into training and test sets for decoding of social, spatial, and combined social–spatial information.

**Supplementary Figure 5.**
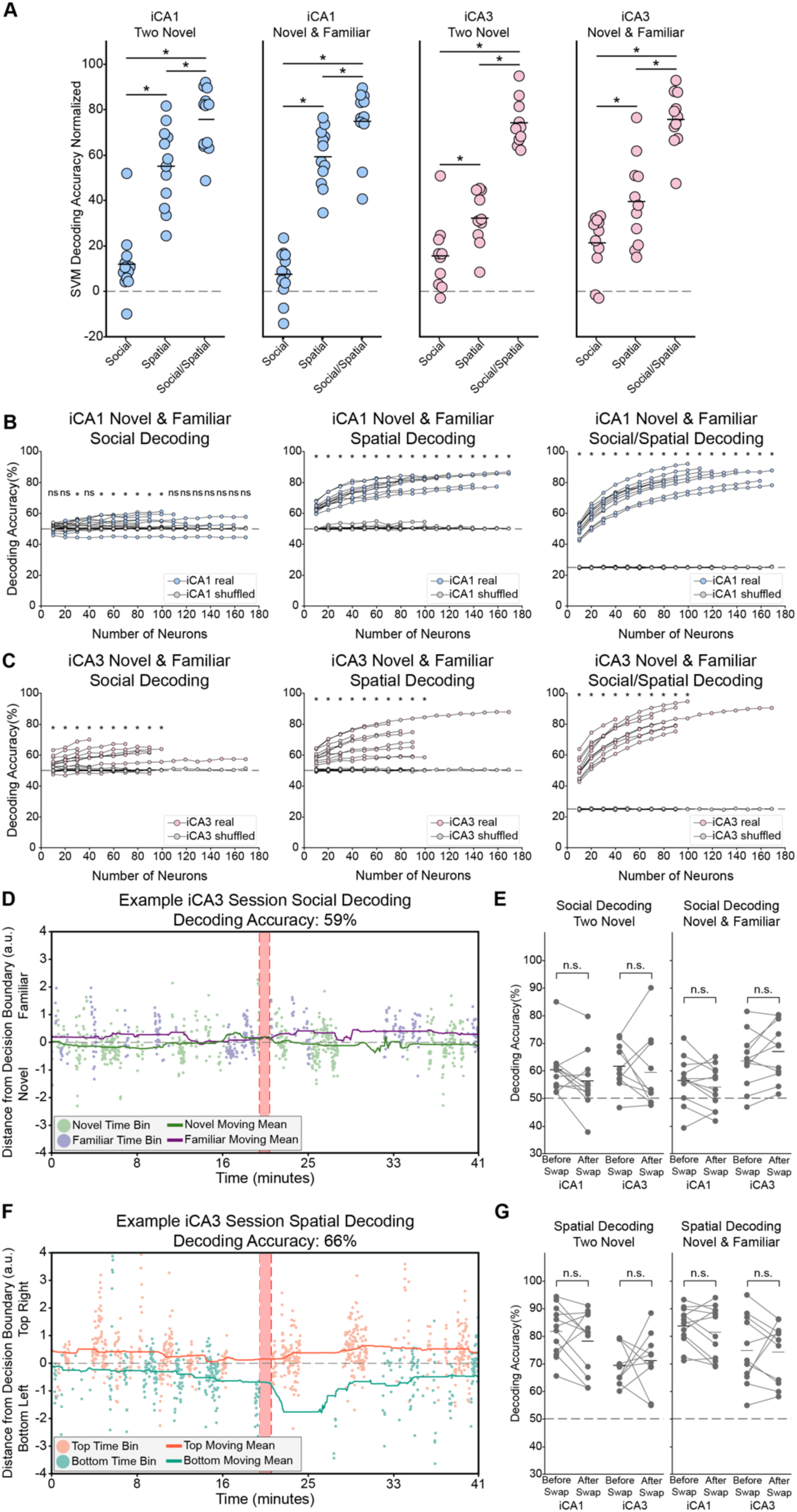
Scaling and Temporal Stability of Social and Spatial Decoding in the SDT. **A)** SVM decoding accuracy from (Fig. 2H-J) was normalized by chance decoding accuracy to allow for direct comparison across decoders. In both the iCA1 and iCA3 cohorts, across both conditions, spatial decoding was significantly higher than social decoding, and social–spatial decoding was significantly higher than both spatial and social decoding, statistics reported in Supplementary Data 1. **B)** Social (left), spatial (center), and social–spatial (right) decoding accuracies in the iCA1 with an increasing number of randomly sampled neurons. Significant effects are marked by asterisks; full test statistics and exact p-values are reported in Supplementary Data 1 **C)** Same as (B), but for the iCA3. **D)** Example session showing the average test set distance from the decision boundary for each time bin when decoding conspecific identity. Novel bins below zero and familiar bins above zero are correctly classified. Importantly, the decoder performs similarly before and after the swap. **E)** Accuracy of time bins before and after the conspecific swap in the two-novel (left) and novel–familiar (right) conditions. Social decoding accuracy did not differ systematically before and after the swap in the two-novel condition in either the iCA1 or the iCA3 (iCA1: paired t-test, t(11) = 2.049, p = 0.066, n = 12; iCA3: paired t-test, t(9) = 0.559, p = 0.590, n = 10). Similarly, there was no significant difference in social decoding accuracy before and after the swap in the novel-familiar condition in either iCA1 and iCA3 (iCA1: paired t-test, t(11) = 1.523, p = 0.156, n = 12; iCA3: paired t-test, t(10) = 1.694, p = 0.121, n = 11). **F)** Same as (D), but for spatial decoding from the same session. Spatial decoding produced stronger separation between classes, with a greater proportion of time bins correctly classified and with greater confidence compared to social decoding. **G)** Same as (E), but for spatial decoders. No significant difference in spatial decoding accuracy was observed before and after the swap in the two-novel condition in either the iCA1 or the iCA3 (iCA1: paired t-test, t(11) = 1.280, p = 0.227, n = 12; iCA3: paired t-test, t(9) = 0.389, p = 0.706, n = 10). Similarly, there was no significant difference in spatial decoding accuracy before and after the swap in the novel-familiar condition in either iCA1 and iCA3 (iCA1: paired t-test, t(11) = 1.321, p = 0.213, n = 12; iCA3: paired t-test, t(10) = 0.1803, p = 0.861, n = 11).

**Supplementary Figure 6.**
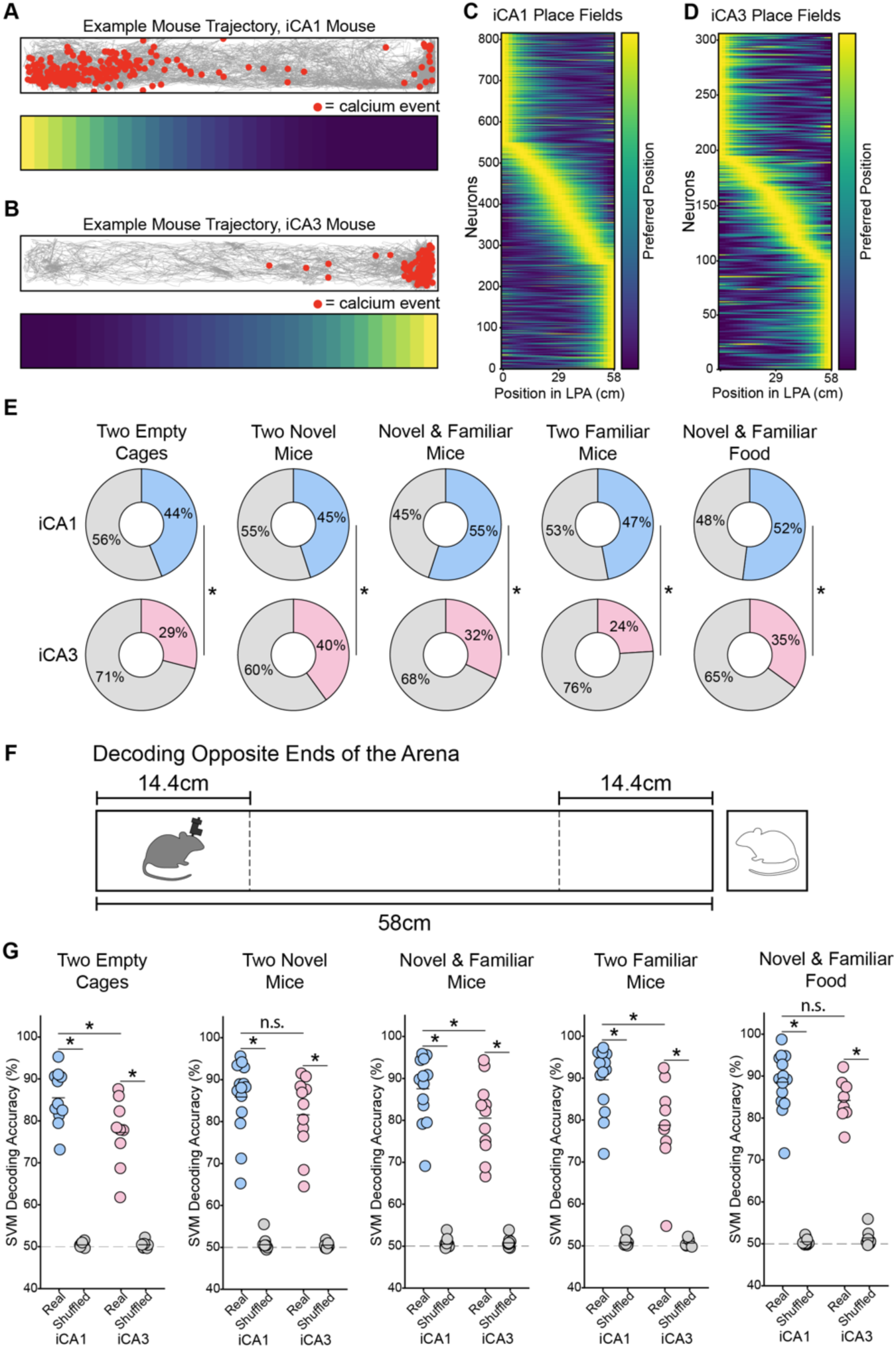
Place Decoding in the Linear Presentation Assay. **A)** Top, trajectory of one imaging mouse (gray) in the LPA overlaid with calcium events (red) from one example iCA1 neuron. Bottom, spatial map of the same neuron. **B)** Same as (A) but for one example iCA3 neuron. **C)** The spatial maps of all significant iCA1 neurons in the novel-familiar conspecific condition. Neurons have been sorted by the position of the place fields. **D)** Same as (C) but for significant iCA3 neurons. **E)** Proportion of significant place cells in iCA1 and iCA3, showing that the iCA1 had consistently more place cells across all conditions. The iCA1 contained a larger proportion of place cells than the iCA3 in the two empty condition (two-proportion z-test, p < 0.001), two-novel conspecifics condition (p = 0.022), the novel-familiar conspecific condition (p < 0.001), two familiar conspecific condition (p < 0.001) and the two-familiar conspecific condition (p < 0.001). **F)** Schematic of the LPA highlighting the 14.4cm region at either end used to decode the position of the imaging mouse. **G)** SVM decoding accuracy in iCA1 and iCA3 when decoding position in the LPA illustrated in (F) from population neural data. Each data point represents the decoding accuracy of one mouse. Both the iCA1 and iCA3 significantly decoded the position of the imaging mouse above chance across all conditions and the iCA1 displayed higher decoding accuracy than the iCA3 in all conditions except for the 2-novel conspecific condition and the novel/familiar food condition. Significant effects are marked by asterisks; full test statistics and exact p-values are reported in Supplementary Data 1.

**Supplementary Figure 7.**
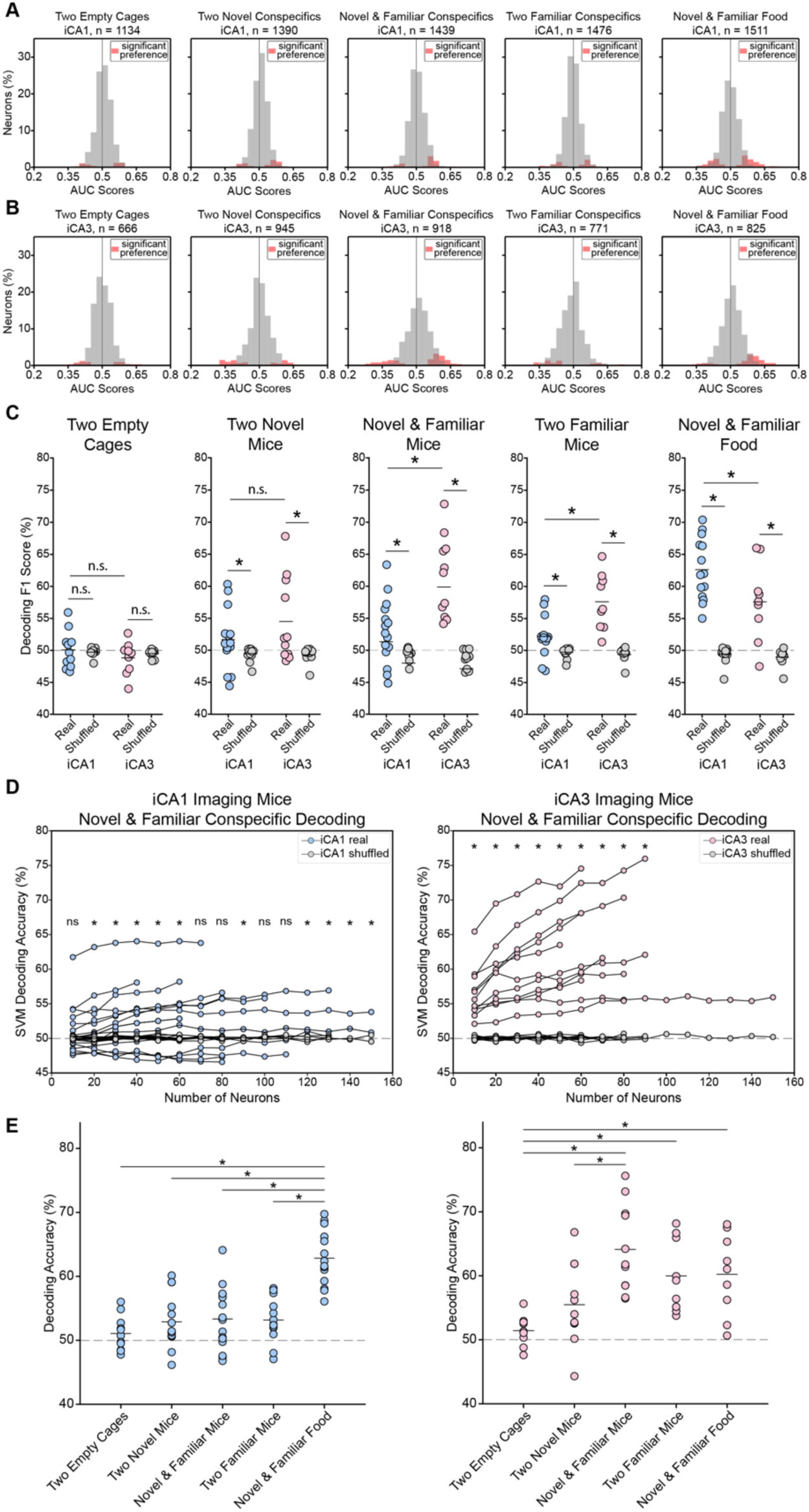
Additional Analyses Supporting LPA Social Encoding and Decoding. **A)** Distribution of AUC values of all iCA1 neurons across LPA behavioral conditions. **B)** Same as (A), but for iCA3 neurons. **C)** SVM decoding accuracy across LPA conditions using F1 scores as the performance metric instead of decoding accuracy; full test statistics are provided in Supplementary Data **D)** Decoding accuracy in the iCA1 (left) and iCA3 (right) in the novel–familiar condition with an increasing number of randomly sampled neurons. **E)** Direct comparison of decoding accuracies between different conditions in the iCA1 (left) and iCA3 (right). All significant results are displayed (hierarchical bootstrap, Bonferroni correction for multiple comparisons; iCA1: 2 empty cages vs. novel/familiar food, Δ = 11.78, p < 0.001, d = 12.306; 2 novel conspecifics vs. novel/familiar food, Δ = 9.943, p < 0.001, d = 8.994; novel/familiar conspecifics vs. novel/familiar food, Δ = 9.495, p < 0.001, d = 8.072; 2 familiar conspecifics vs. novel/familiar food, Δ = 9.67, p < 0.001, d = 9.483; iCA3: 2 empty cages vs. novel/familiar food, Δ = 8.79, p = 0.001, d = 5.786; 2 empty cages vs. 2 familiar conspecifics, Δ = 8.544, p < 0.001, d = 6.225; 2 empty cages vs. novel/familiar conspecifics, Δ = 13.084, p < 0.001, d = 9.218; 2 novel conspecifics vs. novel/familiar conspecifics, Δ = 9.049, p = 0.003, d = 4.917). All statistical comparisons are reported in Supplementary Data 1.

**Supplementary Figure 8.**
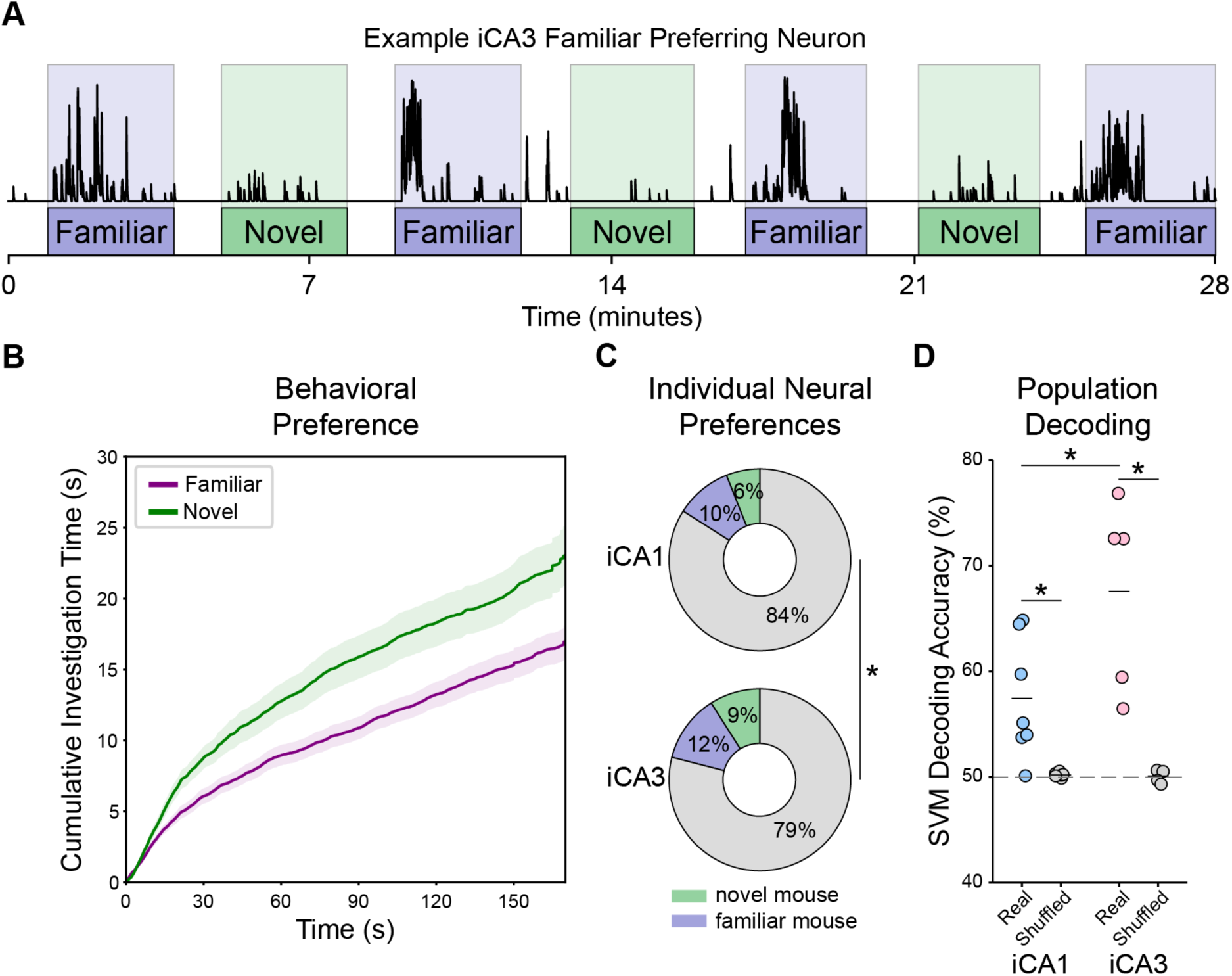
The iCA3 More Strongly Encodes Social Novelty Than the iCA1 in a Free Interaction Test. **(A)** Activity trace of an example iCA3 neuron that preferentially responds to the familiar conspecific (top) and trial structure of the free interaction test (bottom). **(B)** Cumulative time imaging mice investigated the novel and familiar conspecifics, averaged across the first three trials with each conspecific and quantified using SimBA^32^. Imaging mice spent significantly longer investigating the novel conspecific than the familiar conspecific (Wilcoxon signed-rank test: W = 4.00, p = 0.003, n = 12 sessions). **(C)** Proportion of neurons that displayed a preference for one conspecific in the iCA1 (top) and iCA3 (bottom) mice. Proportions were compared between subfields using proportion z-tests (z = 2.212, p = 0.027). **(D)** SVM decoding accuracy in the iCA1 and iCA3 when decoding conspecific identity from population neural data. Both the iCA1 and iCA3 were significantly above chance (iCA1: Δ = 7.23%, p < 0.001, d = 5.052; iCA3: Δ = 17.49%, p < 0.001, d = 6.820) and the iCA3 had significantly higher decoding accuracy than the iCA1 (mean difference = 10.151%, hierarchical bootstrap p = 0.0149, Cohen’s d = 3.506).

**Supplementary Figure 9.**
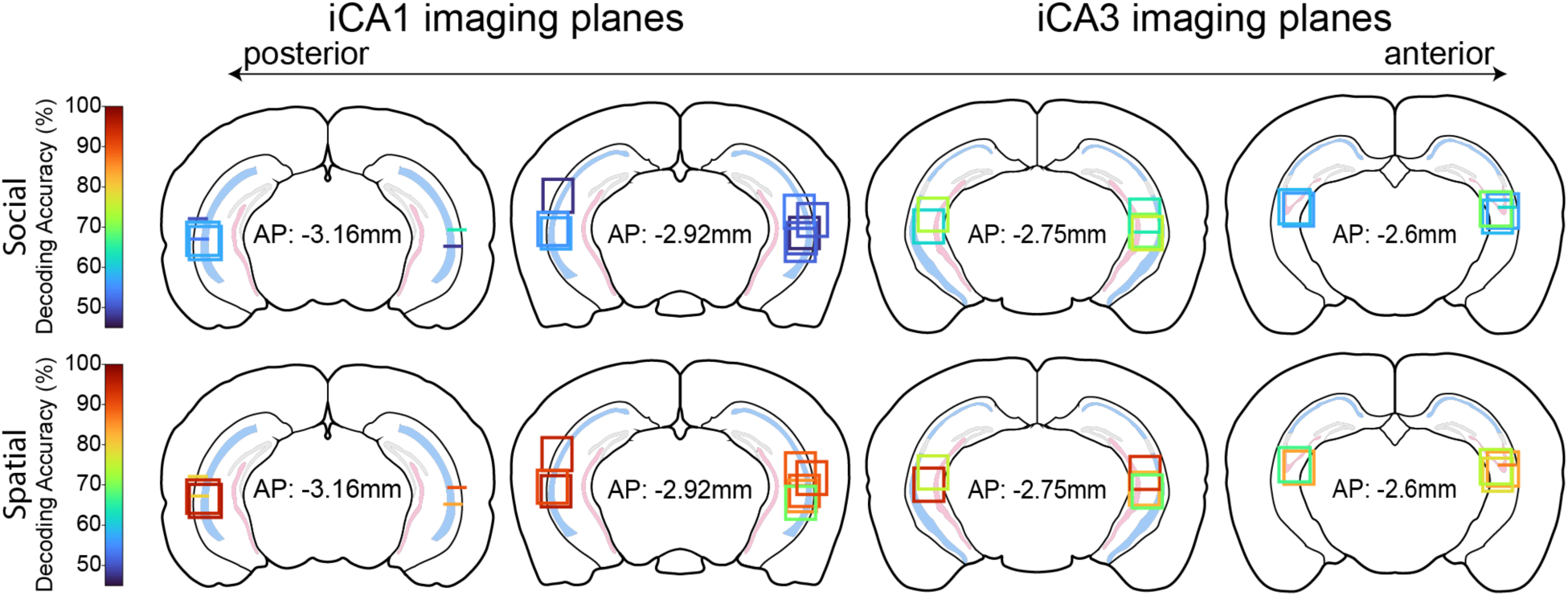
Decoding Accuracy Mapped onto Imaging Planes. SVM decoding accuracy for social (top) and spatial (bottom) information in the novel–familiar conspecific condition, overlaid on reconstructions of imaging planes in the iCA1 and iCA3. Horizontal lines indicate imaging planes obtained with straight lenses, and squares indicate imaging planes obtained with prism lenses. The colors of the lines and squares correspond to the decoding accuracy for social and spatial information.

**Supplementary Figure 10.**
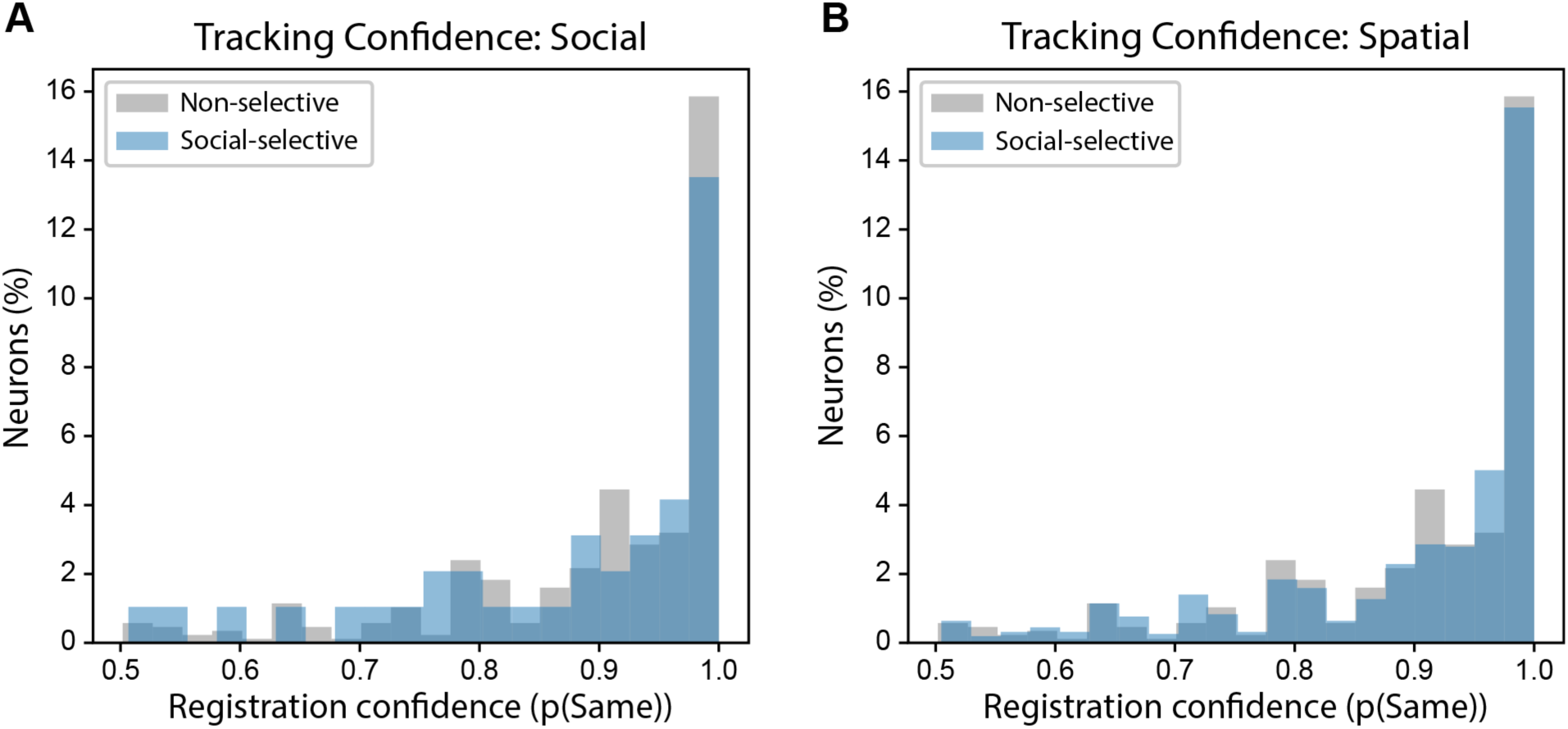
Longitudinally Tracking Confidence Values. (A) Registration probability (P(same), CellReg^77^) for neurons tracked between the two novel/familiar conspecific sessions (Day 5 and Day 7). Neurons displaying a significant social preference in either session were classified as social-selective; neurons showing both social and spatial preferences were excluded from this comparison. There was no significant difference in registration confidence between social-selective and non-selective cells (median P(same): social-selective = 0.939, non-selective = 0.939; two-sample KS test: D = 0.127, p = 0.568). (B) Same as (A), but for neurons displaying a significant spatial preference. There was no significant difference in registration confidence between spatial-selective and non-selective cells (median P(same): spatial-selective = 0.954, non-selective = 0.939; two-sample KS test: D = 0.052, p = 0.557).

### Description of Additional Supplementary Files

**File name: Supplementary Data 1**

*Description: A compendium of all the statistical analyses associated with the figures*.

**File name: Supplementary Movie 1**

*Description: A representative iCA3 imaging animal on the social discrimination task shown with imaging field of view*.

**File name: Supplementary Movie 2**

*Description: A representative iCA3 imaging animal on the linear presentation assay shown with imaging field of view*.

## References

1. Moy, S. S. et al. Sociability and preference for social novelty in five inbred strains: an approach to assess autistic-like behavior in mice. Genes Brain Behav 3, 287–302 (2004).

2. Kogan, J. H., Frankland, P. W. & Silva, A. J. Long-term memory underlying hippocampus-dependent social recognition in mice. Hippocampus 10, 47–56 (2000).

3. Maaswinkel, H., Baars, A. M., Gispen, W. H. & Spruijt, B. M. Roles of the basolateral amygdala and hippocampus in social recognition in rats. Physiol Behav 60, 55–63 (1996).

4. Leroy, F. et al. A circuit from hippocampal CA2 to lateral septum disinhibits social aggression. Nature 564, 213–218 (2018).

5. Meira, T. et al. A hippocampal circuit linking dorsal CA2 to ventral CA1 critical for social memory dynamics. Nat Commun 9, 4163 (2018).

6. Lopez-Rojas, J., de Solis, C. A., Leroy, F., Kandel, E. R. & Siegelbaum, S. A. A direct lateral entorhinal cortex to hippocampal CA2 circuit conveys social information required for social memory. Neuron 110, 1559–1572.e4 (2022).

7. Oliva, A., Fernández-Ruiz, A., Leroy, F. & Siegelbaum, S. A. Hippocampal CA2 sharp-wave ripples reactivate and promote social memory. Nature 587, 264–269 (2020).

8. Boyle, L. M., Posani, L., Irfan, S., Siegelbaum, S. A. & Fusi, S. Tuned geometries of hippocampal representations meet the computational demands of social memory. Neuron 112, 1358–1371.e9 (2024).

9. Hitti, F. L. & Siegelbaum, S. A. The hippocampal CA2 region is essential for social memory. Nature 508, 88–92 (2014).

10. Stevenson, E. L. & Caldwell, H. K. Lesions to the CA2 region of the hippocampus impair social memory in mice. Eur J Neurosci 40, 3294–3301 (2014).

11. Donegan, M. L. et al. Coding of social novelty in the hippocampal CA2 region and its disruption and rescue in a 22q11.2 microdeletion mouse model. Nat Neurosci 23, 1365– 1375 (2020).

12. Okuyama, T., Kitamura, T., Roy, D. S., Itohara, S. & Tonegawa, S. Ventral CA1 neurons store social memory. Science 353, 1536–1541 (2016).

13. Chiang, M.-C., Huang, A. J. Y., Wintzer, M. E., Ohshima, T. & McHugh, T. J. A role for CA3 in social recognition memory. Behav Brain Res 354, 22–30 (2018).

14. Tao, K. et al. Disrupted social memory ensembles in the ventral hippocampus underlie social amnesia in autism-associated Shank3 mutant mice. Mol Psychiatry 27, 2095–2105 (2022).

15. Ishizuka, N., Weber, J. & Amaral, D. G. Organization of intrahippocampal projections originating from CA3 pyramidal cells in the rat. Journal of Comparative Neurology 295, 580– 623 (1990).

16. Sun, Y., Nitz, D. A., Holmes, T. C. & Xu, X. Opposing and Complementary Topographic Connectivity Gradients Revealed by Quantitative Analysis of Canonical and Noncanonical Hippocampal CA1 Inputs. eNeuro 5, (2018).

17. Treves, A. & Rolls, E. T. Computational analysis of the role of the hippocampus in memory. Hippocampus 4, 374–391 (1994).

18. Risold, P. Y. & Swanson, L. W. Connections of the rat lateral septal complex1Published on the World Wide Web on 2 June 1997.1. Brain Research Reviews 24, 115–195 (1997).

19. Gergues, M. M. et al. Circuit and molecular architecture of a ventral hippocampal network. Nat Neurosci 23, 1444–1452 (2020).

20. Isaac, J., Karkare, S. C., Balasubramanian, H. & Murugan, M. Organization of Brainwide Inputs to Discrete Lateral Septum Projection Populations. J. Neurosci. 45, (2025).

21. Yeates, D. C. M. et al. Parallel ventral hippocampus-lateral septum pathways differentially regulate approach-avoidance conflict. Nat Commun 13, 3349 (2022).

22. Eichenbaum, H. On the integration of space, time, and memory. Neuron 95, 1007–1018 (2017).

23. Macbeth, A. H., Edds, J. S. & Young, W. S. Housing conditions and stimulus females: a robust social discrimination task for studying male rodent social recognition. Nat Protoc 4, 1574– 1581 (2009).

24. Schafer, M. & Schiller, D. Navigating Social Space. Neuron 100, 476–489 (2018).

25. Fanselow, M. S. & Dong, H.-W. Are The Dorsal and Ventral Hippocampus functionally distinct structures? Neuron 65, 7 (2010).

26. Jin, S.-W. & Lee, I. Differential encoding of place value between the dorsal and intermediate hippocampus. Current Biology 31, 3053–3072.e5 (2021).

27. Cruz-Sanchez, A. et al. Ventral and Intermediate Hippocampus Are Required for Object-in-Place Recognition Memory in Mice. eNeuro 13, (2026).

28. Jarzebowski, P., Hay, Y. A., Grewe, B. F. & Paulsen, O. Different encoding of reward location in dorsal and intermediate hippocampus. Current Biology 32, 834–841.e5 (2022).

29. Li, K., Koukoutselos, K., Sakaguchi, M. & Ciocchi, S. Distinct ventral hippocampal inhibitory microcircuits regulating anxiety and fear behaviors. Nat Commun 15, 8228 (2024).

30. Xu, C. et al. Distinct Hippocampal Pathways Mediate Dissociable Roles of Context in Memory Retrieval. Cell 167, 961–972.e16 (2016).

31. Watarai, A., Tao, K. & Okuyama, T. Representation of sex-specific social memory in ventral CA1 neurons. Science 389, eadp3814 (2025).

32. Goodwin, N. L. et al. Simple behavioral analysis (SimBA) as a platform for explainable machine learning in behavioral neuroscience. Nat Neurosci 27, 1411–1424 (2024).

33. Kjelstrup, K. B. et al. Finite Scale of Spatial Representation in the Hippocampus. Science 321, 140–143 (2008).

34. Jung, M. W., Wiener, S. I. & McNaughton, B. L. Comparison of spatial firing characteristics of units in dorsal and ventral hippocampus of the rat. J Neurosci 14, 7347–7356 (1994).

35. Kohara, K. et al. Cell type-specific genetic and optogenetic tools reveal novel hippocampal CA2 circuits. Nat Neurosci 17, 269–279 (2014).

36. Mankin, E. A., Diehl, G. W., Sparks, F. T., Leutgeb, S. & Leutgeb, J. K. Hippocampal CA2 activity patterns change over time to a larger extent than between spatial contexts. Neuron 85, 190–201 (2015).

37. Bliss, T. V. P. & Collingridge, G. L. A synaptic model of memory: long-term potentiation in the hippocampus. Nature 361, 31–39 (1993).

38. Malenka, R. C. & Bear, M. F. LTP and LTD: An Embarrassment of Riches. Neuron 44, 5–21 (2004).

39. Nicoll, R. A. & Schmitz, D. Synaptic plasticity at hippocampal mossy fibre synapses. Nat Rev Neurosci 6, 863–876 (2005).

40. Zhao, M., Choi, Y.-S., Obrietan, K. & Dudek, S. M. Synaptic Plasticity (and the Lack Thereof) in Hippocampal CA2 Neurons. J Neurosci 27, 12025–12032 (2007).

41. Simons, S. B., Escobedo, Y., Yasuda, R. & Dudek, S. M. Regional differences in hippocampal calcium handling provide a cellular mechanism for limiting plasticity. Proc Natl Acad Sci U S A 106, 14080–14084 (2009).

42. Pagani, J. H. et al. Role of the vasopressin 1b receptor in rodent aggressive behavior and synaptic plasticity in hippocampal area CA2. Mol Psychiatry 20, 490–499 (2015).

43. Carstens, K. E. & Dudek, S. M. Regulation of synaptic plasticity in hippocampal area CA2. Curr Opin Neurobiol 54, 194–199 (2019).

44. Evans, P. R. et al. RGS14 Restricts Plasticity in Hippocampal CA2 by Limiting Postsynaptic Calcium Signaling. eNeuro 5, (2018).

45. Besnard, A. & Leroy, F. Top-down regulation of motivated behaviors via lateral septum sub-circuits. Mol Psychiatry 27, 3119–3128 (2022).

46. Rashid, M. et al. A ventral hippocampal-lateral septum pathway regulates social novelty preference. eLife 13, (2024).

47. Phillips, M. L., Robinson, H. A. & Pozzo-Miller, L. Ventral hippocampal projections to the medial prefrontal cortex regulate social memory. eLife 8, e44182 (2019).

48. Murugan, M. et al. Combined social and spatial coding in a descending projection from prefrontal cortex. Cell 171, 1663–1677.e16 (2017).

49. Strange, B. A., Witter, M. P., Lein, E. S. & Moser, E. I. Functional organization of the hippocampal longitudinal axis. Nat Rev Neurosci 15, 655–669 (2014).

50. Wang, X. & Zhan, Y. Regulation of Social Recognition Memory in the Hippocampal Circuits. Front. Neural Circuits 16, (2022).

51. Rao, R. P., Heimendahl, M. von, Bahr, V. & Brecht, M. Neuronal Responses to Conspecifics in the Ventral CA1. Cell Reports 27, 3460–3472.e3 (2019).

52. Forli, A. & Yartsev, M. M. Hippocampal representation during collective spatial behaviour in bats. Nature 621, 796–803 (2023).

53. Ray, S. et al. Hippocampal coding of identity, sex, hierarchy, and affiliation in a social group of wild fruit bats. Science 387, eadk9385 (2025).

54. Omer, D. B., Maimon, S. R., Las, L. & Ulanovsky, N. Social place-cells in the bat hippocampus. Science 359, 218–224 (2018).

55. Watarai, A., Tao, K., Wang, M.-Y. & Okuyama, T. Distinct functions of ventral CA1 and dorsal CA2 in social memory. Current Opinion in Neurobiology 68, 29–35 (2021).

56. Wee, R. W. S. et al. Internal-state-dependent control of feeding behavior via hippocampal ghrelin signaling. Neuron 112, 288–305.e7 (2024).

57. Décarie-Spain, L. et al. Ventral hippocampus neurons encode meal-related memory. Nat Commun 16, 4898 (2025).

58. Décarie-Spain, L. et al. Ventral hippocampus-lateral septum circuitry promotes foraging-related memory. Cell Rep 40, 111402 (2022).

59. Liu, Y. et al. A circuit from dorsal hippocampal CA3 to parvafox nucleus mediates chronic social defeat stress–induced deficits in preference for social novelty. Science Advances 8, eabe8828 (2022).

60. Li, M., Jiang, Y.-Q. & Sun, Q. A circuit from basolateral amygdala to hippocampal CA3 regulates social behavior. Curr Biol 35, 4349–4364.e5 (2025).

61. Rolls, E. The mechanisms for pattern completion and pattern separation in the hippocampus. Front. Syst. Neurosci. 7, (2013).

62. Hayashi, Y., Kobayakawa, K. & Kobayakawa, R. The temporal and contextual stability of activity levels in hippocampal CA1 cells. Proceedings of the National Academy of Sciences 120, e2221141120 (2023).

63. Kentros, C. G., Agnihotri, N. T., Streater, S., Hawkins, R. D. & Kandel, E. R. Increased Attention to Spatial Context Increases Both Place Field Stability and Spatial Memory. Neuron 42, 283– 295 (2004).

64. Biane, J. S. et al. Representations of stimulus features in the ventral hippocampus. Neuron 113, 3015–3030.e6 (2025).

65. Mathis, A. et al. DeepLabCut: markerless pose estimation of user-defined body parts with deep learning. Nat Neurosci 21, 1281–1289 (2018).

66. Friard, O. & Gamba, M. BORIS: A free, versatile open-source event-logging software for video/audio coding and live observations. Methods in Ecology and Evolution 7, (2016).

67. Lopes, G. et al. Bonsai: an event-based framework for processing and controlling data streams. Frontiers in Neuroinformatics 9, (2015).

68. Zhou, P. et al. Efficient and accurate extraction of in vivo calcium signals from microendoscopic video data. eLife 7, e28728 (2018).

69. Skaggs, W., McNaughton, B. & Gothard, K. An Information-Theoretic Approach to Deciphering the Hippocampal Code. in Advances in Neural Information Processing Systems vol. 5 (Morgan-Kaufmann, 1992).

70. Pedregosa, F. et al. Scikit-learn: Machine Learning in Python. MACHINE LEARNING IN PYTHON.

71. Remedios, R. et al. Social behaviour shapes hypothalamic neural ensemble representations of conspecific sex. Nature 550, 388–392 (2017).

72. Yang, B., Karigo, T. & Anderson, D. J. TRANSFORMATIONS OF NEURAL REPRESENTATIONS IN A SOCIAL BEHAVIOR NETWORK. Nature 608, 741–749 (2022).

73. Saravanan, V., Berman, G. J. & Sober, S. J. Application of the hierarchical bootstrap to multi-level data in neuroscience. Neuron Behav Data Anal Theory 3, https://nbdt.scholasticahq.com/article/13927-application-of-the-hierarchical-bootstrap-to-multi-level-data-in-neuroscience (2020).

74. Hua, T. et al. General anesthetics activate a potent central pain-suppression circuit in the amygdala. Nat Neurosci 23, 854–868 (2020).

75. Gonzalez, W. G., Zhang, H., Harutyunyan, A. & Lois, C. Persistence of neuronal representations through time and damage in the hippocampus. Science 365, 821–825 (2019).

76. Isaac, J. et al. Sex differences in neural representations of social and nonsocial reward in the medial prefrontal cortex. Nat Commun 15, 8018 (2024).

77. Sheintuch, L. et al. Tracking the Same Neurons across Multiple Days in Ca2+ Imaging Data. Cell Rep 21, 1102–1115 (2017).

